# TetraFuse: A Synergistic Four-Dimensional Dynamic Fusion Framework for Efficient and Robust Medical Image Classification

**DOI:** 10.64898/2026.06.02.729722

**Authors:** Yufei Gao, Jiaqi Li, Jing Xu, Qing Li, Ziyu Li, Guohua Zhao, Yucheng Shi, Xia Wu, Yameng Zhang

## Abstract

Accurate and robust classification of medical pathology images is pivotal for computer-aided diagnosis. However, the deployment of deep learning models in high-throughput clinical screening faces a fundamental challenge: the trade-off between diagnostic accuracy and computational efficiency. Current lightweight architectures, while reducing parameter complexity through grouped convolutions, often lead to cross-channel information isolation and diminished representational capacity. In this paper, we propose **TetraFuse**, a novel framework that systematically integrates features from four complementary domains: space, channel, statistics, and frequency. TetraFuse introduces a novel Cross-Channel Dynamic Aggregation (CCDA) paradigm that reconstructs global channel topology with negligible computational overhead, resolving the inter-group isolation issue. To balance perceptual fidelity and efficiency, we design a stage-aware local enhancement mechanism: Local Variance-Guided Enhancer (LVGE) is employed to filter out shallow-stage background noise, while High-Frequency Boundary Injection (HFBI) reinforces deep-stage pathological contours, preventing spatial over-smoothing. Experimental results on the COVID-19, ISIC 2018, and Kvasir datasets confirm that TetraFuse outperforms state-of-the-art (SOTA) methods. Notably, TetraFuse-Tiny achieves a transformative 91.53% reduction in FLOPs compared to ResNet50; on the Kvasir dataset, it achieved an accuracy of 0.926 and an AUC of 0.994 with only 0.345G FLOPs. By combining high representational power with minimal computational demand, TetraFuse offers a scalable solution for large-scale medical image analysis, especially in resource-constrained clinical environments.

## 1. Introduction

Medical image classification has become a fundamental component of precision medicine and large-scale clinical screening [1; 2]. With the rapid growth of medical imaging data, the ability to accurately and efficiently identify pathological heterogeneity is increasingly paramount for reducing diagnostic workloads and improving clinical decision-making. However, deploying advanced deep learning models in real-world clinical environments remains a formidable challenge. Resource-constrained settings impose strict requirements on computational efficiency and model compactness. Conventional high-performance models frequently rely on deep and parameter-intensive architectures, which, although diagnostically accurate, incur substantial computational overhead and are therefore practically prohibitive for resource-limited clinical applications [3; 4]. In this context, ResNet50 is widely regarded as the clinical standard for medical image analysis due to its robust feature extraction and reliable diagnostic performance, yet its substantial computational demand often serves as the primary barrier for edge-device deployment.

To address these challenges, recent studies have extensively explored lightweight architectures based on grouped and depthwise separable convolutions [5; 6]. While these approaches significantly reduce theoretical computational complexity (e.g., FLOPs), they introduce a critical representational bottleneck: intrinsic information isolation across channels [7; 8]. By partitioning feature channels into independent groups, these methods fragment global feature integration and limit the modeling of inter-channel dependencies. This issue is particularly detrimental in medical image analysis, where diagnostic cues consistently arise from the synergistic interaction of heterogeneous features [9]. For instance, subtle lesion boundaries (high-frequency spatial features) are frequently coupled with chromatic abnormalities across multiple channels. Disrupting such interactions inevitably degrades the model’s diagnostic fidelity. Furthermore, recent benchmarks indicate that simply reducing theoretical FLOPs often results in a significant loss of representational capacity, leading to sub-optimal trade-offs between model size and predictive accuracy [10; 11].

These underlying limitations highlight the urgent need for a novel design paradigm that transcends static and locally constrained feature interactions. To this end, we propose **TetraFuse**, a resource-efficient architecture specifically tailored for robust medical image classification. At the core of TetraFuse is the **Cross-Channel Dynamic Aggregation (CCDA)** paradigm. Unlike conventional grouped convolutions that suffer from rigid information isolation, CCDA reconstructs the global topological integrity of feature channels by facilitating dynamic inter-group interactions with negligible computational overhead [12; 13]. Within this framework, we introduce the **Heteroscale Selective Perception (HSP)** module, which mimics the human visual mechanism of “scanning-then-focusing.” By leveraging a heteroscale-selective block, HSP achieves a multi-scale representations that captures everything from fine-grained local textures to broad global context [14; 15]. Furthermore, the **Local Feature Aggregation (LFA)** module is integrated, utilizing grouped dynamic kernels to achieve fine-grained spatial extraction while maintaining an exceptionally sparse parameter footprint [16].

Recognizing that medical imaging data possesses distinct statistical properties across network depths, TetraFuse incorporates a domain-aware feature enhancement strategy. In the shallow stages, we propose the **Local Variance-Guided Excitation (LVGE)** module. Moving beyond traditional first-order attention, LVGE utilizes second-order statistical variance to distinguish between smooth physiological tissues and heterogeneous pathological lesions, effectively purifying early-stage features from background noise [17; 18]. Conversely, in the deep stages, where sequential operations act as low-pass filters [19], the **High-Frequency Boundary Injection (HFBI)** module is deployed. HFBI employs a frequency-spatial decoupling strategy to explicitly reinstate high-frequency residual signals into the deep semantic features, thereby counteracting spatial over-smoothing and preserving the sharp structural boundaries critical for lesion contour identification [20; 21].

The main contributions of this work are summarized as follows:

- **A Resource-Efficient Architectural Paradigm:** We propose TetraFuse, a highly scalable network that reconciles the long-standing conflict between diagnostic representational power and computational parsimony.
- **Global-Local Dynamic Interaction:** We introduce the CCDA and LFA paradigms, which collectively bridge the information gap in lightweight models. By synergizing with the HSP module, the network achieves multi-scale adaptive perception, facilitating a seamless transition from global context modeling to microscopic detail extraction.
- **Statistical and Frequency-Aware Enhancement:** We propose two specialized enhancers: LVGE, which leverages second-order statistical heterogeneity for early-stage noise suppression, and HFBI, which utilizes frequency-domain decoupling to safeguard structural integrity against deep stage semantic over smoothing.
- **State-of-the-Art Performance-Efficiency Balance:**

Extensive evaluations demonstrate TetraFuse achieves superior accuracy and AUC across multiple pathological datasets. Notably, the model delivers top-tier performance with a minimal FLOPs footprint of only 0.345G (91.53% reduction compared to ResNet50). This comparison demonstrates that TetraFuse can achieve heavyweight diagnostic fidelity at an ultralightweight computational cost, proving its high suitability for resource-constrained clinical deployment.

## 2. Related Work

This section reviews key advancements that contribute to the evolution of efficient medical image classification models, focusing on four major aspects: lightweight network architectures, efficient computational block designs, attention mechanisms, and frequency-aware feature enhancement strategies.

### 2.1. Lightweight Architectures

Advancements in convolutional neural networks (CNNs) have revolutionized AI-assisted integration in medical imaging [1; 2]. While classical architectures such as ResNet [22] and DenseNet [23] achieve exceptional diagnostic accuracy, their immense parameter counts and heavy computational burdens render them impractical for deployment in resource-constrained clinical environments [3; 24]. To address this, lightweight paradigms such as MobileNet [6; 8] and ShuffleNet [5] utilized depthwise separable convolutions to dramatically reduce theoretical FLOPs. However, this efficiency frequently compromises the model’s ability to extract rich, cross-channel diagnostic cues. Recently, Vision Transformers (ViTs) [25] and Mamba-based architectures [26; 21] have been introduced into the medical domain to capture long-range global dependencies [27; 28]. Nevertheless, ViTs suffer from high computational overhead and a lack of inductive bias, while standard lightweight CNNs struggle with representational bottlenecks [29]. The evolution of these architectures highlights an urgent need for models that can balance state-of-the-art diagnostic performance with minimal computational footprints, a core motivation behind the proposed TetraFuse. Specifically, the DAFNet [30] achieved significant progress in pathology image classification by focusing on salient lesion regions. However, it primarily optimizes for accuracy, leaving a gap in the exploration of model deployment efficiency for high-throughput clinical screening, which TetraFuse aims to address.

### 2.2. Efficient Computational Block Design

The architectural design of computational blocks is pivotal to network performance, defining the fundamental feature extraction capabilities. To mitigate the computational cost of dense convolutions, modern lightweight blocks extensively adopt grouped convolutions and channel shuffling mechanisms [7]. While efficient, rigidly partitioning channels inevitably leads to inter-group information isolation, disrupting the holistic topology of multi-dimensional features [9]. To improve representational expressiveness, Li et al. [31] proposed Selective Kernel Networks (SKNet) featuring multiple branches with different kernel sizes, and Yu et al. [14] integrated large-kernel designs to emulate multiscale visual perception. Similarly, Chen et al. [32] introduced dynamic convolutions to adaptively adjust aggregation weights, while Ding et al. [16] utilized structural re-parameterization to maintain training robustness. Despite these innovations, existing blocks typically fail to simultaneously solve the channel isolation problem and maintain lightweight characteristics. Building upon these foundations, our proposed CCDA block and LFA module reconstruct global channel topology and perform multiscale dynamic aggregation without imposing significant computational overhead. Unlike the feature fusion strategy in DAFNet [30] that may introduce high-dimensional redundancy and computational overhead, TetraFuse employs the CCDA module to achieve cross-channel alignment with negligible parameters. This allows for the reconstruction of global topology which was often neglected in previous attention-based fusion models like DAFNet.

### 2.3. Attention Mechanism

Beyond static computational blocks, attention mechanisms have emerged as a pivotal paradigm, empowering models to focus dynamically on salient pathological features. Hu et al. [13] compressed feature maps via global average pooling to implement channel attention (SE), whereas Wang et al. [12] introduced a lightweight 1D convolution (ECA) to facilitate local cross-channel interactions efficiently. Woo et al. [33] combined both spatial and channel attention to refine feature maps sequentially. In the context of medical imaging, attention is crucial for distinguishing lesions from complex backgrounds [17; 18]. However, conventional attention modules predominantly rely on first-order statistics (e.g., mean activation values), which frequently falsely highlight bright artifacts or homogeneous background noise rather than genuine pathological heterogeneity. Recognizing this limitation, our Local Variance-Guided Excitation (LVGE) module diverges from traditional approaches by leveraging second-order statistical variance to explicitly purify shallow-stage textures. While DAFNet [30] effectively captures long-range dependencies through a dual-attention mechanism (spatial and channel), it remains limited to a two-dimensional perspective. In contrast, our TetraFuse extends this paradigm by integrating statistical and frequency domain information, forming a more robust four-dimensional dynamic fusion.

### 2.4. Frequency-Aware Enhancement

In deep neural networks, sequential convolution and pooling operations inherently act as low-pass filters, progressively over-smoothing feature representations [19]. While advantageous for extracting abstract global semantics, this phenomenon severely degrades high-frequency structural details, such as subtle lesion boundaries, which are critical for medical image classification [34]. To counteract this, Huo et al. [15] proposed a hierarchical multi-scale fusion network to maintain feature diversity across different abstraction levels. Bi et al. [20] utilized a cross-level consistency strategy to preserve structural integrity under complex degradation. These studies collectively demonstrate that safeguarding high-frequency components significantly enhances clinical diagnostic robustness. Inspired by this, TetraFuse integrates the High-Frequency Boundary Injection (HFBI) module. By explicitly decoupling frequency and spatial domains, HFBI effectively reinjects high-frequency residual signals into deep semantic features, ensuring that the network maintains microscopic precision alongside global contextual understanding.

## 3. Proposed Method

The proposed lightweight TetraFuse is a novel architecture specifically tailored for robust medical image classification. By synergistically integrating Local Variance-Guided Excitation (LVGE), Cross-Channel Dynamic Aggregation (CCDA), and High-Frequency Boundary Injection (HFBI), the TetraFuse effectively reconciles the trade-off between perceptual fidelity and computational efficiency [2; 10].

This hierarchical design enables a seamless transition from early-stage texture purification to late-stage structural reinforcement, substantially augmenting the representational power of lightweight CNNs in the presence of intense pathological heterogeneity and background noise. In the following sections, we will provide a comprehensive description of the mathematical formulations and the architectural details of these components.

### 3.1. TetraFuse Architecture

The proposed TetraFuse is shown in Fig. 1. The overall structure of the network comprises a MSC-Stem layer, four CCDAPaths, four DDS layers and a classifier. Detailed descriptions of these components are provided in the following sections.

**Figure 1.**
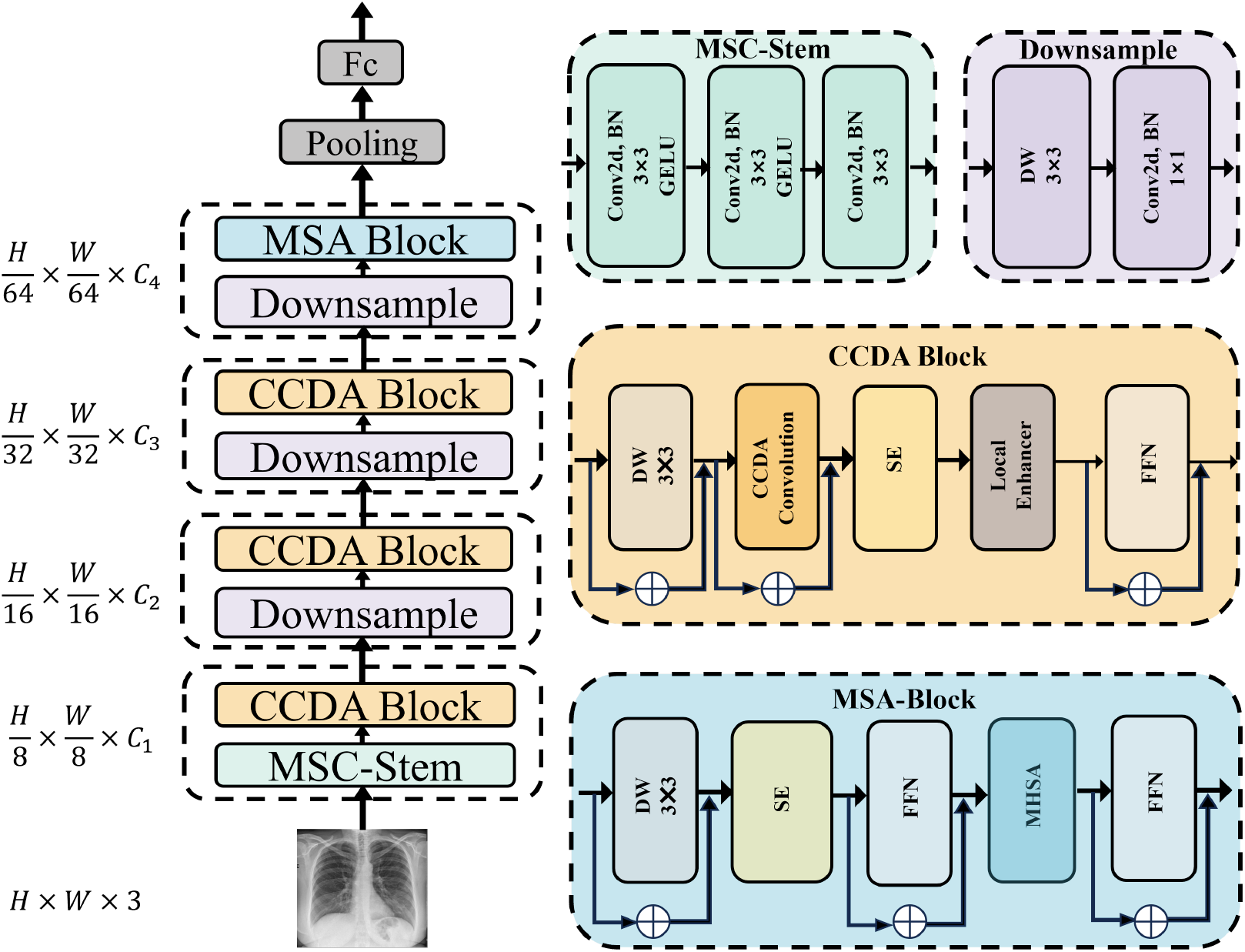
The architecture of the TetraFuse and details of each component. TetraFuse has four stages with 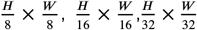, and 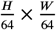 resolutions respectively, where *H* and *W* denote the width and height of the input image. *C* represents the channel dimension. The norm layer and nonlinearity are omitted for simplicity.

#### 3.1.1. MSC-Stem

As illustrated in Fig. 1, the proposed Multi-Stage Convolutional Stem (MSC-Stem) progressively maps the input image into a high-dimensional feature space through a sequence of strided convolution operations. In contrast to standard single-layer patch embedding [25], MSC-Stem utilizes multiple layers with small-sized convolutional kernels to enhance the perception granularity of initial features [4; 15].

The operations are formally defined in Eqs. (1)–(3):

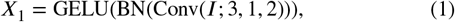

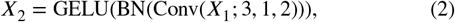

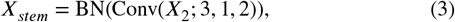

where Conv(*X*; *k, p, s*) denotes the convolution operation on input *X* with kernel size *k*, padding *p*, and stride *s*. Each layer is integrated with Batch Normalization (BN) and GELU activation [35]. *I* represents the input image, *X*_1_ and *X*_2_ are intermediate feature maps, and *X*_*stem*_ signifies the final output produced by the stem network.

#### 3.1.2. Multi CCDAPaths of TetraFuse

Following the initial feature extraction by MSC-stem, the resulting feature maps are passed through four connecting pathways with some differences, termed CCDAPaths. CCDAPath1 and CCDAPath2 consist of some CCDABlocks with LVGE module, followed by a Decoupled Down-Sampling (DDS) module. Within each path, the features are processed through the CCDABlocks sequentially, with the outputs filtered and then downsampled by the corresponding DDS Module. CCDAPath3 utilizes the same structure but the LVGE module is replaced by the HFBI module. CCDAPath4 adopts a similar structure, but the most CCDABlock is replaced by a Multi-Head Self-Attention (MSA) block [36]. This design allows CCDAPath4 to integrate global semantic features, providing more comprehensive information for subsequent classification [21].

To adapt more flexibly to datasets of varying scales, we have developed three model variants tailored to specific scales. Table 1 details the corresponding hyperparameter configurations for the different CCDAPath counts.

**Table 1.** Hyperparameters of the proposed model variants.

| Model | Path1 | Path2 | Path3 | Path4 |
| --- | --- | --- | --- | --- |
| TetraFuse_Tiny | 0 | 1 | 4 | 0 |
| TetraFuse_Small | 1 | 2 | 4 | 0 |
| TetraFuse_Base | 2 | 3 | 4 | 2 |

#### 3.1.3. DDS Module

To facilitate hierarchical representation while minimizing the loss of critical pathological information during resolution reduction, we design the Decoupled Down-Sampling (DDS) module at stage transitions. The DDS layer decouples spatial filtering from channel projection to preserve topological integrity [8; 11], as described in Eqs. (4)–(5):

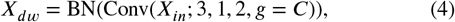

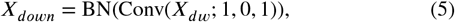

where *g* = *C* represents a depth-wise convolution (*C* is the number of input channels) [6]. This design ensures efficient feature propagation and maintains a lightweight parameter footprint during the down-sampling process.

#### 3.1.4. Multi-head Self-Attention (MSA) Block

The MSA block is strategically integrated into the final stage of TetraFuse to facilitate global dependency modeling. Unlike the convolutional layers in earlier stages that focus on local textures, the MSA block enables the network to aggregate information across the entire spatial domain [25]. The forward pass of the MSA module is formulated as in Eqs. (6)–(10):

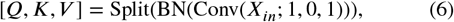

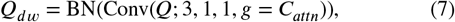

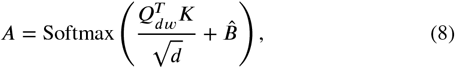

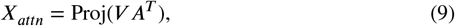

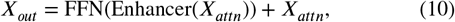

where *Q, K*, and *V* denote the Query, Key, and Value matrices, respectively. To incorporate inductive bias, a depthwise convolution is applied to *Q* to generate *Q*_*dw*_ [16; 37], while 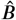 provides a learnable relative position encoding. The HFBI enhancer subsequently refines the attended features to ensure that high-frequency boundary details are preserved alongside the global context.

#### 3.1.5. Diagnostic Classifier

The classification head is designed to project high-level semantic features into the pathological diagnostic space. After performing Global Average Pooling (GAP), a BN-Linear structure is employed to stabilize the latent distribution and generate final predictions, as defined in Eqs. (11)–(12):

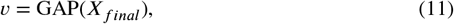

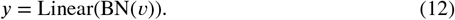

For scenarios involving knowledge distillation [38], the classifier concurrently generates a primary prediction vector *y* and a distillation vector *y*_*dist*_, thereby achieving an optimal balance between model efficiency and diagnostic accuracy.

### 3.2. CCDA Convolution

The details of CCDA Convolution is shown in Fig. 2. In recent years, lightweight visual architectures have widely adopted grouped convolution or depthwise separable convolution as alternatives to dense standard convolution [6; 5]. To emulate the functionality of the human vision system, LSNet employed grouped convolutions to achieve lightweight spatial aggregation. However, this approach inevitably resulted in information isolation between channels [7]. During the feature extraction process, channels are rigidly partitioned into mutually independent subspaces, resulting in the mutual information flow between groups being completely severed. In the domain of medical image processing, such isolation is detrimental [9]. For instance, in the context of multi-label lesion classification tasks, the diagnosis of lesions frequently relies on the joint distribution of cross-channel features. Specifically, irregular textural boundaries, which correspond to high-frequency spatial information, are typically highly coupled with abnormal pigmentation, which corresponds to cross-channel color features.

**Figure 2.**
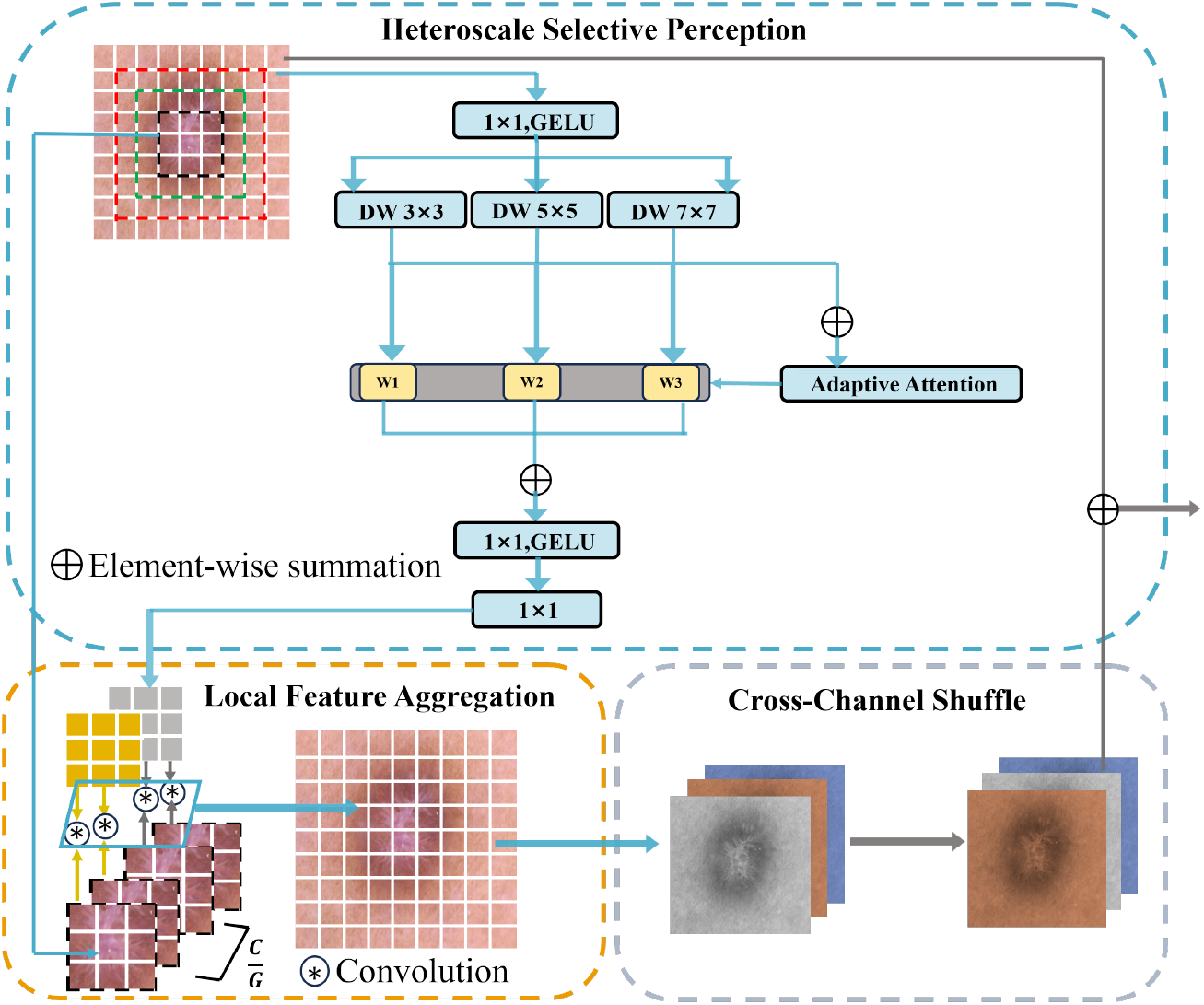
The architecture of the CCDA Convolution and details of each component.

To overcome this bottleneck, a departure from traditional static linear interactions has been effected, and a novel spatial-channel decoupling paradigm has been proposed: Cross-Channel Dynamic Aggregation (CCDA). CCDA aims to reconstruct the topological integrity of global channels at the network’s lower layers without imposing additional computational overhead, thereby achieving deep semantic alignment of multi-dimensional features.To further clarify the structural innovations of CCDA and its boundaries with existing dynamic mechanisms, we provide a comparative analysis with Selective Kernel Networks (SKNet) [31] and Dynamic Convolution [32]. As summarized in Table 2, CCDA distinguishes itself through a synergistic cross-channel alignment paradigm rather than simple kernel-level selection or fusion.

**Table 2.**
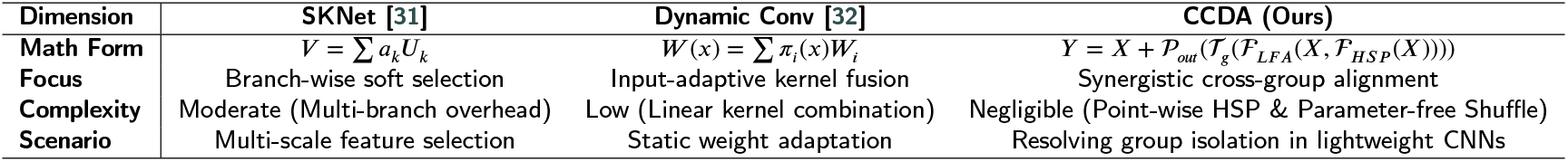
Differentiation analysis between CCDA and existing dynamic convolution methods.

The proposed CCDA Convolution introduces an effective computational architecture that synergistically integrates three key components: the Heteroscale Selective Perception (HSP) module, the Local Feature Aggregation (LFA) module, and Cross-Channel Shuffle with Mixing. The entire process of the CCDA convolution, from input **X** to output **Y**, can be expressed by Eq. (13):

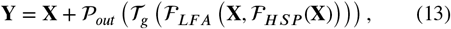

where ℱ _*HSP*_ (·) denotes the heteroscale perception mapping that generates the dynamic weight tensor **A**_*dyn*_, and ℱ _*LFA*_ (·) represents the group-wise dynamic aggregation guided by these weights. 𝒯_*g*_ signifies the channel shuffle operator used to break the inter-group isolation [7], while 𝒫_*out*_ refers to the final point-wise mixer initialized with a zero-residual strategy [39] to ensure training stability. By integrating these components, CCDA Convolution effectively achieves high-order non-linear modeling of both local textures and global channel correlations. The whole architecture of CCDA Convoution is illustrated in Fig. 2.

#### 3.2.1. Cross-Channel Shuffle with Mixing

To alleviate the inherent limitation of grouped feature aggregation, we introduce a channel shuffling operation after the LFA-based intra-group dynamic aggregation. Although LFA effectively enhances spatial feature representation within each group, it inevitably restricts inter-group information exchange [7].

Specifically, given an input feature map **X** ∈ ℝ^*B*×*C*×*H*×*W*^, the operation first reshapes it into 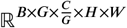, where *G* denotes the number of groups. It then performs a transpose between the group dimension and the intra-group channel dimension, followed by flattening back to the original shape. This reshape–transpose–flatten process effectively implements a deterministic channel permutation, redistributing channels from the same group across different groups. This parameter-free operation introduces negligible computational overhead while effectively breaking the group-wise isolation, enabling sufficient inter-group feature interaction.

More importantly, the shuffled features become well-mixed along the channel dimension, providing a more informative and balanced input for the subsequent cross-channel linear fusion. After that, to enable effective cross-channel feature interaction, we employ a lightweight cross-channel linear mixer implemented as a 1 × 1 convolution followed by batch normalization. This design follows a reparameterization paradigm [16], balancing training stability and inference efficiency.

#### 3.2.2. HSP Module

Heteroscale Selective Perception (HSP) adopts the design of a heteroscale-selective block. In complex medical images, such as dermatological lesions or histopathological sections, the distribution of diagnostic information exhibits pronounced cross-scale heterogeneity [31]. Traditional single-scale convolutional kernels frequently encounter difficulties in capturing this multi-scale information within a single pathway. The HSP module simulates the dynamic perception mechanism of “scanning first, then focusing” in the human visual system, achieving multi-scale representations from local details to global context [14].

Given a visual feature map **X** ∈ ℝ^*H*×*W* ×*C*^, we initially utilize a point-wise convolution 𝒫_1_(⋅) to project the tokens into a lower channel dimension (*C*/2 by default) as in Eq. (14):

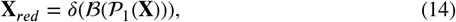

where ℬ and *δ* represent Batch Normalization and GELU activation, respectively. We then employ heteroscale depth-wise convolution (DW) with a set of kernel sizes 𝒦 = {3, 5, 7} to capture different spatial contextual fields. For each *k* ∈ 𝒦, the spatial information is extracted via Eq. (15):

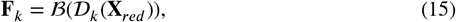

where 𝒟_*k*_ denotes the DW convolution with kernel size *k* × *k*. We then leverage a Selective Decision Unit (SDU) to calculate the contribution weight for the three branches. The SDU first aggregates the multiscale features **U** = ∑_*k* ∈ 𝒦_ **F**_*k*_ and extracts the global context *s* = 𝒢 (**U**) through global average pooling 𝒢 (⋅) [13]. The attention weight vector ***ω*** = [*ω*_1_, *ω*_2_, *ω*_3_]^*T*^ is generated via a bottleneck architecture [12] in Eq. (16):

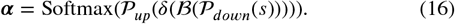

The fused feature **X**_*fused*_ achieves dual adaptability in both spatial and scale dimensions, formulated as Eq. (17):

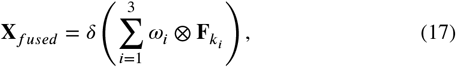

where ⊗ denotes the element-wise Hadamard product. Finally, a point-wise convolution is utilized to generate the context-dependent perception weights **A**_*dyn*_ ∈ ℝ^*H*×*W* ×*D*^ for the subsequent aggregation in Eq. (18):

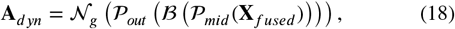

where 𝒩_*g*_ is the Group Normalization layer.

#### 3.2.3. LFA Module

Local Feature Aggregation (LFA) employs grouped dynamic convolutions to achieve fine-grained local feature extraction. Given the feature map **X** ∈ ℝ^*H*×*W* ×*C*^ passed from the HSP module, we divide its channels into *G* groups. Each group contains *C*/*G* channels, and channels within the same group share identical aggregation weights.

For each center feature token *x*_*i*_ at a spatial location, we receive the multi-scale adaptive weights **A**_*dyn*_ from the HSP module. We define a mapping operator ℛ (⋅) that reshapes the weight vector into a structured grouped dynamic kernel as defined in Eq. (19):

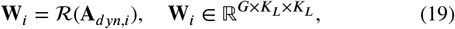

where *K*_*L*_ × *K*_*L*_ represents the microscopic receptive field. The aggregation response for the *c*-th channel of token *x*_*i*_ belonging to the *g*-th group is computed as Eq. (20):

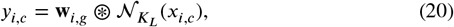

where ⊛ denotes the dynamic grouping convolution operator [32] and 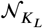 represents the local neighborhood. By sharing weights within each group, the LFA module reduces the parameter count to 1/*G* relative to standard dynamic convolutions, ensuring high sensitivity to local pathological details without incurring prohibitive computational over-head.

### 3.3. Local Enhancer

To refine spatial fidelity and compensate for the potential information loss during feature aggregation, a Local Feature Enhancer (Ψ) is integrated into the block. This component adaptively employs two specialized modules, LVGE and HFBI, to address the distinct representational requirements of different stages:

- **Local Variance Guided Enhancer (LVGE):** The details of LVGE is shown in Fig. 3. In the shallow stages, LVGE is utilized to purify local textures. It leverages local variance to suppress noise and emphasize finegrained anatomical details [17; 18].
- **High-Frequency Boundary Injector (HFBI):** The details of HFBI is shown in Fig. 4. In the deep stages, HFBI is deployed to reinforce structural integrity. By capturing high-frequency components, it injects sharp boundary information back into the feature maps, effectively counteracting the smoothing effect of global self-attention [20; 21].

**Figure 3.**
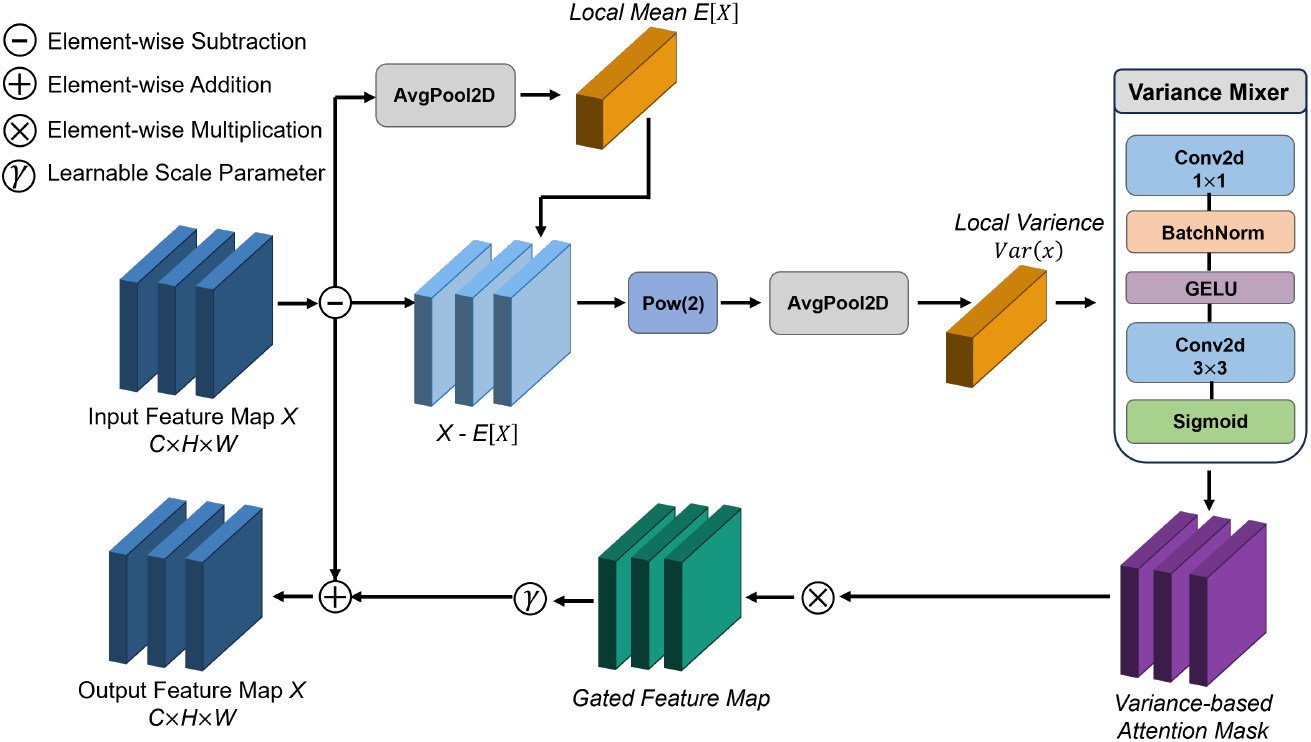
The architecture of the Local Variance-Guided Excitation (LVGE) module. The module explicitly captures irregular lesion textures by computing the local variance Var(*X*) (orange block). This statistical prior is then transformed into a Variance-based Attention Mask (purple block) to adaptively highlight potential lesion regions while suppressing smooth background noise.

**Figure 4.**
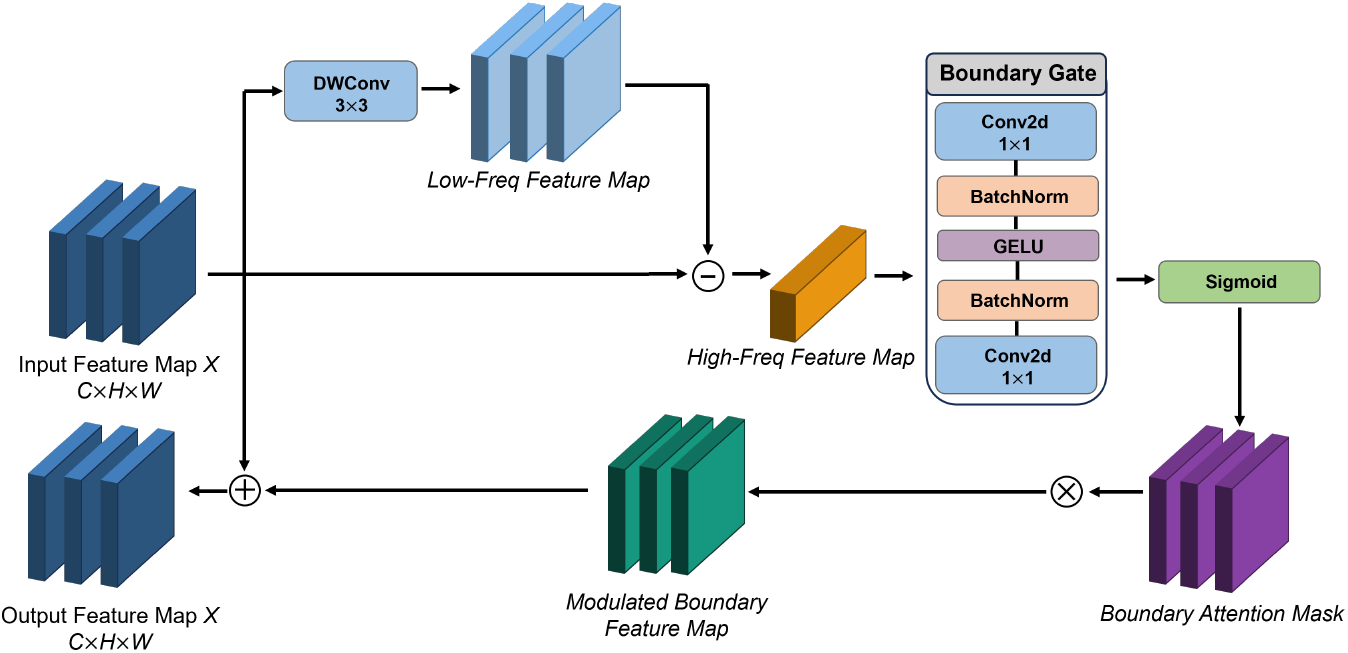
The schematic of the High-Frequency Boundary Injector (HFBI). HFBI decouples high-frequency cues (orange block) via a differencing operation between the input and its smoothed low-frequency counterpart. A Boundary Attention Mask (purple block) is subsequently generated to precisely inject structural details into the primary feature stream.

The operation of the Local Enhancer is formulated in Eqs. (21)–(23):

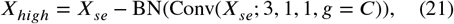

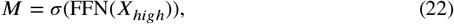

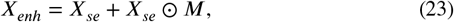

where *X*_*high*_ represents the high-frequency residual extracted via depth-wise convolution (*g* = *C*), and *M* denotes the learnable enhancement mask. This mechanism ensures that TetraFuse maintains a robust balance between global semantic understanding and local pathological precision.

#### 3.3.1. LVGE Module

In the shallow stages of the network, high-resolution feature maps are frequently susceptible to interference from medical background noises. Conventional spatial attention mechanisms typically rely on absolute activation values (first-order statistics) [13], which often falsely highlight bright artifacts. We hypothesize that healthy physiological tissues exhibit smooth, low-variance distributions, whereas pathological lesions demonstrate intense structural heterogeneity. Based on this prior, the LVGE module is proposed to suppress homogeneous noise [17].

Given an input feature **X** ∈ ℝ^*H*×*W* ×*C*^, the LVGE module first estimates the local statistical heterogeneity. Let ℰ(⋅) denote the local mean operator implemented via a sliding window. The local spatial variance **V** is computed as shown in Eq. (24):

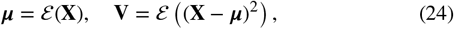

where **V** ∈ ℝ^*H*×*W* ×*C*^ represents the second-order statistical distribution. To transform the raw variance map into a robust guidance mask, we employ a variance-mixer ℱ _*mix*_ defined in Eq. (25):

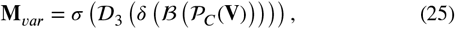

where *σ* denotes the Sigmoid function, and 𝒟_3_ is a 3 × 3 depth-wise convolution used to smooth the variance distribution. The utilization of second-order statistics (local variance) is predicated on the observation that in shallow feature maps, stochastic noise typically exhibits high-frequency isotropic variance, whereas structural edges possess anisotropic local gradients. By calculating the local variance, LVGE can effectively distinguish between these semantic signals and background artifacts in the early stages of the network. Finally, the output feature **Y** is modulated through a residual excitation mechanism as in Eq. (26):

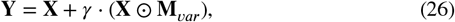

where *γ* is a learnable scaling parameter initialized to zero.

#### 3.3.2. HFBI Module

As the network depth increases, sequential convolutional and pooling operations act as strong low-pass filters, frequently leading to the over-smoothing of features [19]. The High-Frequency Boundary Injector (HFBI) is designed to explicitly extract and reinject high-frequency residuals through a frequency-spatial decoupling strategy.

Given a deep feature map **X** ∈ ℝ^*H*×*W* ×*C*^, we first generate a low-frequency approximation 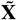 via a depth-wise convolution 𝒟_3_ and batch normalization ℬ in Eq. (27):

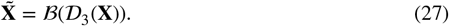

A pure high-frequency residual **H**_*freq*_, encapsulating abrupt textural changes, is derived via subtraction [20] in Eq. (28):

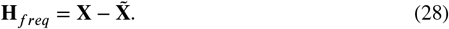

To transform the raw high-frequency signals into boundary guidance, we employ a bottleneck-structured gating mechanism ℱ_*gate*_ formulated as Eq. (29):

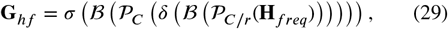

where *r* denotes the reduction ratio. The final output **Y** is obtained by injecting the boundary-sensitive features back into the original representation via Eq. (30):

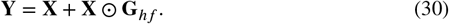

By enhancing the response of pixels located at intensity discontinuities, HFBI guarantees that abstract semantic features remain highly sensitive to subtle lesion contours, effectively counteracting the blurring effects of deep pooling.

## 4. Experiment

### 4.1. Implementation details

To rigorously assess the effectiveness and robustness of TetraFuse, we conducted different experiments on three medical image classification datasets, each of them is a public benchmark. TetraFuse consistently outperformed state-of-the-art and baseline models with diverse architectures across all benchmarks. In the following sections, the experiments were structured around four key components: (1) dataset description and evaluation metric description, (2) experiment details, (3) comparison experiments, and (4) ablation analysis. Fig. 5 presents samples from each category in each dataset.

**Figure 5.**
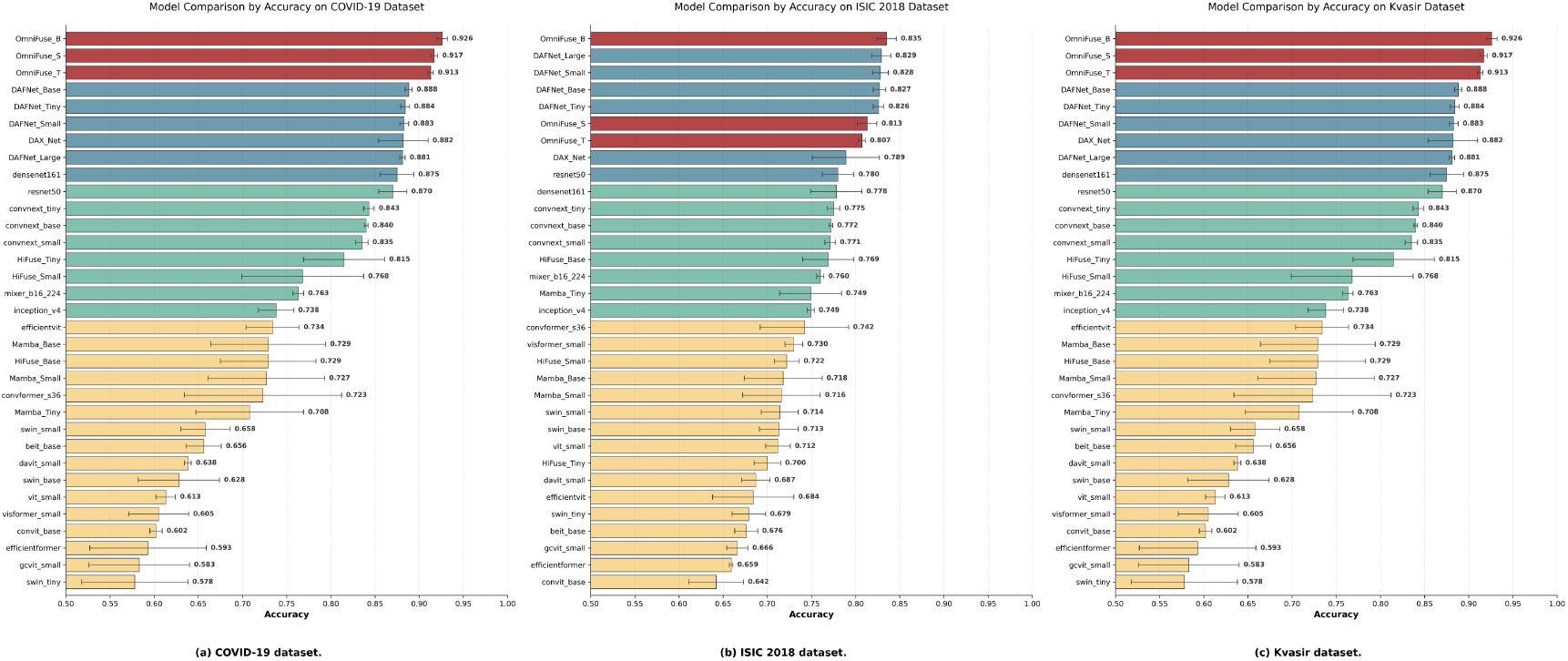
Comparison of classification accuracy of different models on three medical imaging datasets.

### 4.2. Datasets description

#### COVID-19 chest X-ray dataset [52; 53]

This dataset comprises three categories of images, including 1525 COVID-19 positive chest X-ray images, 1525 chest X-ray images depicting other lung diseases, and 1525 normal chest X-ray images. The image resolutions ranged form 224 × 224 to 5623 × 4757 pixels, with the majority having a resolution of 1024 × 1024 pixels. Some of these images contain watermarks or other marks.

#### ISIC2018 (International Skin Imaging Collaboration) dataset [54]

We utilize the ISIC2018 dataset for our experiments,which is specifically designed for the task of skin lesion diagnosis. It serves as a standard benchmark in medical image analysis. The disease classfication task (Task 3) comprises of 11,720 high-resolution images from seven different categories, including melanoma (malignant), melanocytic nevus (benign), basal cell carcinoma (malignant), actinic keratosis (premalignant), benign keratosis (benign), dermatofibroma (benign), and vascular lesion (benign). The original image size in the dataset is 650 × 450 pixels.

#### Kvasir gastroenterology dataset [55]

This dataset comprises gastrointestinal (GI) tract images collected using endoscopic equipment at the Vestre Viken Health Trust in Norway. It comprises 8,000 high-quality images (ranging from 720 × 576 to 1920 × 1072 pixels) categorized into eight distinct classes, each containing 1,000 samples. These classes encompass:

- **Anatomical landmarks:** cecum, ileum, and pylorus;
- **Pathological findings:** esophagitis, polyps, and ulcerative colitis;
- **Clinical procedures:** dyed-resection margins and dyed-polyps.

The balanced distribution and significant intra-class morphological variations make Kvasir a robust benchmark for validating the generalization and discriminative power of deep learning architectures in clinical endoscopic analysis

### 4.3. Evaluation metrics

We utilized a comprehensive set of evaluation metrics for our classification tasks, including Accuracy (ACC), F1 Score, Precision, Recall, Mathews Correlation Coefficient (MCC) [56], Cohen’s Kappa [57], and Area Under the Curve (AUC) [58]. We utilized the confusion matrix which is a standard tool for evaluating the classification performance by comparing predicted labels with ground truth annotations. We also presented the ROC Curve on three datasets to show the AUC results. In addition, we also measures the parameters, and FLOPs of each model.

(1) Experimental Setup: We resized all images to 224 × 224 to ensure the same input size. All experiments were processing with Python 3.8 and PyTorch 2.1.2. For images processing, we adopted basic data augmentation techniques, such as random cropping and normalization.To ensure the statistical robustness and generalizability of TetraFuse, we adopted a rigorous data partitioning and evaluation strategy. The complete dataset was initially partitioned into three independent subsets: a training set (70%), a validation set (20%), and a strictly held-out test set (10%).

To mitigate potential bias and optimize model parameters, a five-fold cross-validation protocol was executed on the combined pool of the training and validation data (the 90% portion). Under this scheme, the dataset was divided into five distinct folds, where each fold served as the validation set once while the remaining four folds were used for training and hyperparameter tuning. The final performance metrics, including Accuracy (Acc) and F1-score, are reported as the average results across the five independent folds evaluated on the fixed, unseen 10% test set. This hierarchical approach ensures that the reported diagnostic efficacy is not the result of over-fitting to a specific data split. Given the inherent class imbalance in the ISIC 2018 dataset (e.g., the ratio of Melanoma to Melanocytic Nevus is approximately 1:7), we implemented two primary strategies to mitigate bias. First, a Weighted Cross-Entropy (WCE) loss function was adopted during training, assigning higher penalty weights to the minority classes to prevent the model from converging toward the majority class. Second, we prioritized the F1-score and Area Under the ROC Curve (AUC) as our primary evaluation metrics rather than simple accuracy, as they provide a more objective assessment of the model’s diagnostic performance on imbalanced medical data. In addition, the AdamW optimizer with a weight decay coefficient of 0.01 was used. Training was performed with a batch size of 32 for 300 training epochs on an NVIDIA RTX 4090 GPU with 24 GB of memory. An early stopping criterion was applied, ending the training if the validation performance did not improve for consecutive 20 epochs. The initial learning rate was set to 1e-4 and decayed step by step to a minimun 1e-6. We performed all comparative experiments under the same enviroment with the same hyperparameter settings. The experimental configuration is provided in Table 3.

**Table 3.**
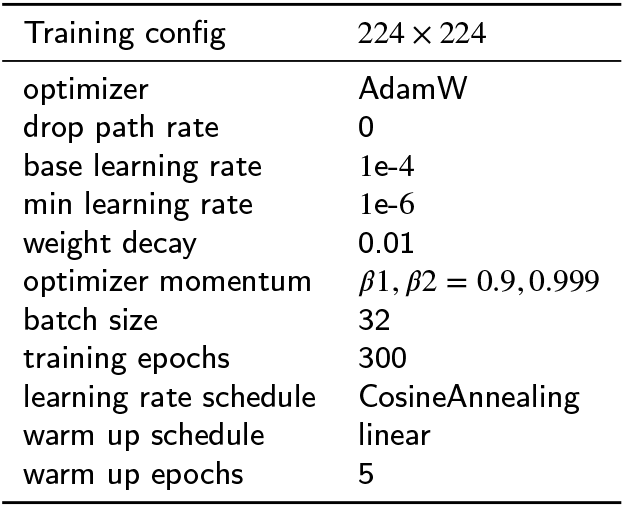
Experimental setting.

(2) Comparative Models: To evaluate the performance of the proposed model, we processed comparative experiments with state-of-the-art CNNs, transformer-based models, and hybrid models that combine CNNs and transformers, all of that are widely used in computer vision and medical image analysis. The models we used included: Beit [40], Gcvit [41], Convformer [47], Convit [42], Efficientvit [43], Efficientformer [11], Swin_transformer [45], Visformer [46], Davit [44], DenseNet [23], ResNet [49], ConvNeXt [50], Inception [4], and Mixer [48], as well as medical image-specific models such as HiFuse [15], DAX_Net [51], MedMamba [21] and the newest state-of-the-art DAFNet [30].

### 4.4. Experiment results

In this section, we report the results on the three benchmark datasets.

#### 4.4.1. COVID-19 dataset

We fully compared the proposed model with other state-of-the-art models in recent years. The experimental results were summarized in Table 4. Top three performances were highlighted using consistent notation: the best result was underlined, the second best was double-underlined, and the third best was wavy underlined. As shown in Table 4, TetraFuse outperformed all other models across most evaluation metrics. Specially, TetraFuse_Base achieved the highest performance across multiple metrics, ranking first in accuracy (ACC: 0.956 ± 0.003), F1 score (0.956 ± 0.005), MCC (0.935 ± 0.008), Kappa (0.934 ± 0.008), AUC (0.990 ± 0.005) and specificity (0.979 ± 0.004). Notably, most CNN-based hybrid models performed better than Transformer-based models on this dataset. DAFNet_Base and ConvNeXt demonstrated stronger performance, due to convolution kernels force the model to focus on neighbouring pixels. In medical imaging, pneumonia is typically characterized by localized changes in texture. The architecture of the CNN is designed with the assumption that neighbouring pixels are strongly correlated, which enables it to converge rapidly even when data is limited. The superior results of TetraFuse can be attributed to the CCDA Convolution and LVGE enhancer, which enhance the local feature and focus on details. CCDA Convolution utilizes convolutional kernels of varying sizes to fully fuse multi-scale features whilst minimizing the loss of local information. By leveraging local variance to rapidly localize the textural heterogeneity of lesions, LVGE enhancer employs attention masks in the early stages to suppress artifacts, thereby preventing the propagation of irrelevant information into deeper layers. Fig. 5 (a) presents a bar chart comparing ACC, illustrating that TetraFuse consistently achieved the highest performance.

**Table 4.** Model performance comparison on COVID-19 dataset [65,66] with top-3 ranking markers: <u>single underline</u> (1st), <u>double underline</u> (2nd), <u>wavy underline</u> (3rd). All ranked values are bolded.

| Model | Acc | F1 | MCC | Kappa | AUC | Specificity |
| --- | --- | --- | --- | --- | --- | --- |
| beit_base [40] | 0.702 ± 0.065 | 0.699 ± 0.064 | 0.561 ± 0.096 | 0.553 ± 0.098 | 0.858 ± 0.035 | 0.851 ± 0.033 |
| gcvit_small [41] | 0.808 ± 0.038 | 0.808 ± 0.066 | 0.717 ± 0.005 | 0.712 ± 0.006 | 0.931 ± 0.002 | 0.904 ± 0.019 |
| convit_base [42] | 0.806 ± 0.101 | 0.806 ± 0.042 | 0.714 ± 0.065 | 0.709 ± 0.052 | 0.936 ± 0.011 | 0.903 ± 0.021 |
| efficientvit [43] | 0.847 ± 0.052 | 0.848 ± 0.051 | 0.774 ± 0.078 | 0.770 ± 0.078 | 0.958 ± 0.024 | 0.923 ± 0.026 |
| efficientformer [11] | 0.769 ± 0.059 | 0.770 ± 0.070 | 0.670 ± 0.086 | 0.653 ± 0.089 | 0.910 ± 0.050 | 0.884 ± 0.030 |
| davit_small [44] | 0.853 ± 0.002 | 0.853 ± 0.066 | 0.783 ± 0.001 | 0.779 ± 0.003 | 0.941 ± 0.005 | 0.926 ± 0.001 |
| swin_tiny [45] | 0.792 ± 0.064 | 0.785 ± 0.106 | 0.704 ± 0.040 | 0.688 ± 0.096 | 0.929 ± 0.029 | 0.896 ± 0.032 |
| swin_small [45] | 0.863 ± 0.023 | 0.863 ± 0.018 | 0.795 ± 0.025 | 0.794 ± 0.025 | 0.959 ± 0.007 | 0.931 ± 0.012 |
| swin_base [45] | 0.796 ± 0.023 | 0.797 ± 0.038 | 0.699 ± 0.040 | 0.694 ± 0.034 | 0.914 ± 0.013 | 0.898 ± 0.011 |
| visformer_small [46] | 0.899 ± 0.021 | 0.900 ± 0.019 | 0.854 ± 0.017 | 0.849 ± 0.021 | 0.971 ± 0.001 | 0.950 ± 0.010 |
| vit_small [25] | 0.799 ± 0.027 | 0.796 ± 0.036 | 0.709 ± 0.063 | 0.698 ± 0.090 | 0.917 ± 0.056 | 0.899 ± 0.024 |
| convformer_s36 [47] | 0.920 ± 0.011 | 0.921 ± 0.010 | 0.883 ± 0.016 | 0.880 ± 0.016 | 0.980 ± 0.001 | 0.960 ± 0.005 |
| mixer_b16_224 [48] | 0.898 ± 0.005 | 0.899 ± 0.005 | 0.849 ± 0.008 | 0.847 ± 0.008 | 0.971 ± 0.002 | 0.949 ± 0.003 |
| densenet161 [23] | 0.924 ± 0.014 | 0.924 ± 0.017 | 0.891 ± 0.017 | 0.886 ± 0.015 | 0.974 ± 0.010 | 0.962 ± 0.012 |
| resnet50 [49] | 0.913 ± 0.009 | 0.913 ± 0.019 | 0.873 ± 0.023 | 0.869 ± 0.023 | 0.949 ± 0.015 | 0.956 ± 0.005 |
| convnext_tiny [50] | 0.933 ± 0.007 | 0.933 ± 0.007 | 0.902 ± 0.010 | 0.899 ± 0.011 | 0.984 ± 0.001 | 0.966 ± 0.004 |
| convnext_small [50] | 0.929 ± 0.009 | 0.929 ± 0.010 | 0.894 ± 0.015 | 0.893 ± 0.014 | 0.984 ± 0.002 | 0.964 ± 0.005 |
| convnext_base [50] | 0.931 ± 0.014 | 0.931 ± 0.014 | 0.898 ± 0.021 | 0.896 ± 0.021 | 0.985 ± 0.002 | 0.966 ± 0.007 |
| inception_v4 [4] | 0.900 ± 0.033 | 0.900 ± 0.035 | 0.854 ± 0.048 | 0.850 ± 0.050 | 0.969 ± 0.010 | 0.950 ± 0.017 |
| HiFuse_Tiny [15] | 0.905 ± 0.041 | 0.905 ± 0.041 | 0.860 ± 0.060 | 0.857 ± 0.061 | 0.971 ± 0.017 | 0.953 ± 0.020 |
| HiFuse_Small [15] | 0.907 ± 0.023 | 0.908 ± 0.023 | 0.864 ± 0.034 | 0.861 ± 0.033 | 0.973 ± 0.007 | 0.954 ± 0.011 |
| HiFuse_Base [15] | 0.929 ± 0.011 | 0.928 ± 0.011 | 0.895 ± 0.016 | 0.893 ± 0.016 | 0.980 ± 0.005 | 0.964 ± 0.005 |
| DAX_Net [51] | 0.918 ± 0.020 | 0.919 ± 0.020 | 0.880 ± 0.030 | 0.878 ± 0.030 | 0.976 ± 0.006 | 0.959 ± 0.010 |
| Mamba_Tiny [21] | 0.862 ± 0.048 | 0.863 ± 0.049 | 0.798 ± 0.071 | 0.793 ± 0.071 | 0.961 ± 0.013 | 0.931 ± 0.024 |
| Mamba_Small [21] | 0.888 ± 0.010 | 0.889 ± 0.010 | 0.836 ± 0.014 | 0.832 ± 0.015 | 0.965 ± 0.004 | 0.944 ± 0.005 |
| Mamba_Base [21] | 0.848 ± 0.031 | 0.847 ± 0.032 | 0.777 ± 0.042 | 0.771 ± 0.046 | 0.954 ± 0.009 | 0.923 ± 0.015 |
| DAFNet_Tiny [30] | 0.938 ± 0.005 | 0.938 ± 0.005 | 0.909 ± 0.007 | 0.906 ± 0.003 | 0.981 ± 0.003 | 0.969 ± 0.002 |
| DAFNet_Small [30] | 0.942 ± 0.003 | 0.942 ± 0.003 | 0.915 ± 0.006 | 0.913 ± 0.006 | <u>0.987 ± 0.002</u> | 0.971 ± 0.002 |
| DAFNet_Base [30] | <u>0.944 ± 0.005</u> | <u>0.944 ± 0.005</u> | <u>0.917 ± 0.008</u> | <u>0.915 ± 0.008</u> | <u>0.987 ± 0.001</u> | <u>0.972 ± 0.002</u> |
| DAFNet_Large [30] | 0.940 ± 0.001 | 0.940 ± 0.002 | 0.912 ± 0.003 | 0.909 ± 0.002 | 0.984 ± 0.003 | 0.970 ± 0.001 |
| TetraFuse_Tiny | 0.941 ± 0.005 | 0.940 ± 0.005 | 0.911 ± 0.007 | 0.911 ± 0.003 | 0.982 ± 0.003 | 0.970 ± 0.002 |
| TetraFuse_Small | <u>0.949 ± 0.004</u> | <u>0.949 ± 0.002</u> | <u>0.924 ± 0.006</u> | <u>0.924 ± 0.004</u> | 0.986 ± 0.003 | <u>0.975 ± 0.005</u> |
| TetraFuse_Base | <u>0.956 ± 0.003</u> | <u>0.956 ± 0.005</u> | <u>0.935 ± 0.008</u> | <u>0.934 ± 0.008</u> | <u>0.990 ± 0.005</u> | <u>0.979 ± 0.004</u> |

#### 4.4.2. ISIC 2018 dataset

In order to provide more comprehensive validation of the superiority of TetraFuse, we conducted comparative experiments on ISIC 2018 dataset using the same experimental conditions as that used on the COVID-19 dataset. We present the detailed results across all the evaluation metrics in Table 5, and the visual result is shown in Fig. 5 (b). On the ISIC 2018 dataset, while the Base version of TetraFuse achieves state-of-the-art performance, outperforming all compared models, achieving top scores in ACC (0.835 ± 0.011), F1 score (0.722 ± 0.011), MCC (0.710 ± 0.011), Kappa (0.710 ± 0.011), AUC (0.966 ± 0.011) and specificity (0.961 ± 0.002), all with lower variance, demonstrating strong efficiency and stability, the Tiny and Small variants exhibit a more balanced profile between diagnostic accuracy and computational overhead. It is worth noting that for the ISIC 2018 dataset, the ACC and F1-score of TetraFuse-Tiny and TetraFuse-Small are slightly lower than those of the DAFNet series. This can be attributed to the prioritized design for clinical deployment efficiency in TetraFuse. Specifically, DAFNet relies on high-dimensional feature redundancy which enhances metrics on specific datasets but at the cost of significantly higher FLOPs. In contrast, TetraFuse utilizes the CCDA module to achieve cross-channel synergy. This optimization ensures that TetraFuse maintains a superior performance-efficiency curve, making it more suitable for high-throughput screening in resource-constrained medical environments. Even the more computationally efficient variants, TetraFuse_Tiny and TetraFuse_Small, consistently outperformed larger, more complex models such as Hi-Fuse_Base and DAX_Net. This trend demonstrates that architectural design of TetraFuse excels at capturing discriminative features without requiring excessive parameter over-head. TetraFuse maintains a steady and narrow error margin across diverse metrics like MCC and Kappa, underscoring its robust generalization capability. By delivering both high peak performance and exceptional predictive stability, TetraFuse proves to be a more dependable framework for realworld medical image classification than its less consistent counterparts.

**Table 5.** Model performance comparison on ISIC 2018 dataset [67] with top-3 ranking markers: single underline (1st), double underline (2nd), <u>wavy underline</u> (3rd). All ranked values are bolded.

| Model | Acc | F1 | MCC | Kappa | AUC | Specificity |
| --- | --- | --- | --- | --- | --- | --- |
| beit_base [40] | 0.676 ± 0.013 | 0.190 ± 0.055 | 0.189 ± 0.095 | 0.137 ± 0.097 | 0.843 ± 0.038 | 0.874 ± 0.013 |
| gcvit_small [41] | 0.666 ± 0.012 | 0.146 ± 0.061 | 0.074 ± 0.131 | 0.067 ± 0.128 | 0.598 ± 0.137 | 0.867 ± 0.019 |
| convit_base [42] | 0.642 ± 0.031 | 0.145 ± 0.027 | 0.114 ± 0.096 | 0.094 ± 0.082 | 0.754 ± 0.032 | 0.871 ± 0.013 |
| efficientvit [43] | 0.684 ± 0.046 | 0.203 ± 0.179 | 0.111 ± 0.221 | 0.110 ± 0.220 | 0.643 ± 0.148 | 0.873 ± 0.032 |
| efficientformer [11] | 0.659 ± 0.002 | 0.130 ± 0.021 | 0.064 ± 0.070 | 0.043 ± 0.061 | 0.711 ± 0.026 | 0.863 ± 0.008 |
| davit_small [44] | 0.687 ± 0.016 | 0.287 ± 0.062 | 0.305 ± 0.054 | 0.271 ± 0.062 | 0.854 ± 0.037 | 0.895 ± 0.010 |
| swin_tiny [45] | 0.679 ± 0.019 | 0.202 ± 0.085 | 0.197 ± 0.164 | 0.180 ± 0.157 | 0.788 ± 0.094 | 0.884 ± 0.024 |
| swin_small [45] | 0.714 ± 0.021 | 0.320 ± 0.100 | 0.391 ± 0.072 | 0.369 ± 0.083 | 0.886 ± 0.026 | 0.911 ± 0.014 |
| swin_base [45] | 0.713 ± 0.022 | 0.307 ± 0.080 | 0.398 ± 0.075 | 0.379 ± 0.087 | 0.890 ± 0.022 | 0.914 ± 0.014 |
| visformer_small [46] | 0.730 ± 0.010 | 0.399 ± 0.032 | 0.436 ± 0.029 | 0.419 ± 0.034 | 0.907 ± 0.003 | 0.917 ± 0.006 |
| vit_small [25] | 0.712 ± 0.014 | 0.376 ± 0.022 | 0.414 ± 0.009 | 0.402 ± 0.008 | 0.899 ± 0.010 | 0.917 ± 0.003 |
| convformer_s36 [47] | 0.742 ± 0.050 | 0.455 ± 0.161 | 0.485 ± 0.127 | 0.475 ± 0.135 | 0.910 ± 0.037 | 0.929 ± 0.020 |
| mixer_b16_224 [48] | 0.760 ± 0.004 | 0.520 ± 0.012 | 0.524 ± 0.014 | 0.519 ± 0.016 | 0.920 ± 0.004 | 0.934 ± 0.004 |
| densenet161 [23] | 0.778 ± 0.029 | 0.582 ± 0.079 | 0.570 ± 0.061 | 0.568 ± 0.062 | 0.934 ± 0.016 | 0.942 ± 0.009 |
| resnet50 [49] | 0.780 ± 0.018 | 0.581 ± 0.048 | 0.572 ± 0.035 | 0.570 ± 0.034 | 0.934 ± 0.010 | 0.941 ± 0.004 |
| convnext_tiny [50] | 0.775 ± 0.007 | 0.571 ± 0.024 | 0.559 ± 0.022 | 0.555 ± 0.024 | 0.932 ± 0.003 | 0.939 ± 0.005 |
| convnext_small [50] | 0.771 ± 0.006 | 0.566 ± 0.012 | 0.553 ± 0.011 | 0.551 ± 0.011 | 0.930 ± 0.003 | 0.940 ± 0.002 |
| convnext_base [50] | 0.772 ± 0.002 | 0.565 ± 0.011 | 0.556 ± 0.001 | 0.554 ± 0.003 | 0.931 ± 0.004 | 0.940 ± 0.001 |
| inception_v4 [4] | 0.749 ± 0.004 | 0.459 ± 0.011 | 0.485 ± 0.007 | 0.473 ± 0.008 | 0.912 ± 0.003 | 0.924 ± 0.002 |
| HiFuse_Tiny [15] | 0.700 ± 0.015 | 0.325 ± 0.032 | 0.369 ± 0.034 | 0.346 ± 0.040 | 0.873 ± 0.017 | 0.908 ± 0.009 |
| HiFuse_Small [15] | 0.722 ± 0.014 | 0.382 ± 0.053 | 0.419 ± 0.042 | 0.402 ± 0.049 | 0.898 ± 0.018 | 0.915 ± 0.008 |
| HiFuse_Base [15] | 0.769 ± 0.029 | 0.541 ± 0.116 | 0.552 ± 0.069 | 0.547 ± 0.073 | 0.931 ± 0.015 | 0.939 ± 0.012 |
| DAX_Net [51] | 0.789 ± 0.038 | 0.570 ± 0.137 | 0.587 ± 0.084 | 0.584 ± 0.087 | 0.938 ± 0.023 | 0.943 ± 0.012 |
| Mamba_Tiny [21] | 0.749 ± 0.035 | 0.488 ± 0.109 | 0.501 ± 0.083 | 0.497 ± 0.085 | 0.913 ± 0.029 | 0.932 ± 0.013 |
| Mamba_Small [21] | 0.716 ± 0.044 | 0.390 ± 0.157 | 0.423 ± 0.117 | 0.409 ± 0.130 | 0.889 ± 0.039 | 0.920 ± 0.021 |
| Mamba_Base [21] | 0.718 ± 0.044 | 0.362 ± 0.150 | 0.390 ± 0.134 | 0.372 ± 0.146 | 0.870 ± 0.063 | 0.910 ± 0.023 |
| DAFNet_Tiny [30] | 0.826 ± 0.006 | 0.689 ± 0.022 | 0.665 ± 0.014 | 0.661 ± 0.016 | 0.951 ± 0.007 | 0.953 ± 0.005 |
| DAFNet_Small [30] | <u>0.828 ± 0.009</u> | 0.713 ± 0.023 | 0.689 ± 0.025 | <u>0.688 ± 0.026</u> | 0.954 ± 0.004 | <u>0.958 ± 0.005</u> |
| DAFNet_Base [30] | 0.827 ± 0.007 | <u>0.718 ± 0.011</u> | <u>0.690 ± 0.011</u> | <u>0.688 ± 0.011</u> | <u>0.955 ± 0.008</u> | <u>0.958 ± 0.002</u> |
| DAFNet_Large [30] | <u>0.829 ± 0.011</u> | <u>0.719 ± 0.012</u> | <u>0.696 ± 0.008</u> | <u>0.694 ± 0.009</u> | <u>0.957 ± 0.012</u> | <u>0.960 ± 0.004</u> |
| TetraFuse_Tiny | 0.807 ± 0.004 | 0.679 ± 0.022 | 0.663 ± 0.014 | 0.659 ± 0.016 | 0.935 ± 0.002 | 0.951 ± 0.005 |
| TetraFuse_Small | 0.813 ± 0.011 | 0.703 ± 0.023 | 0.683 ± 0.025 | 0.682 ± 0.026 | 0.953 ± 0.004 | 0.957 ± 0.005 |
| TetraFuse_Base | <u>0.835 ± 0.011</u> | <u>0.722 ± 0.011</u> | <u>0.710 ± 0.011</u> | <u>0.710 ± 0.011</u> | <u>0.966 ± 0.011</u> | <u>0.961 ± 0.002</u> |

#### 4.4.3. Kvasir dataset

Further experiment was conducted on the Kvasir dataset to test the generalization capability of TetraFuse. The whole evaluation results are shown in Table 6. TetraFuse significantly outperformed other models in the classification task. Obviously, TetraFuse_Base achieved the best results in all key metrics: ACC (0.926 ± 0.006), F1 score (0.925 ± 0.005), MCC (0.915 ± 0.004), Kappa (0.915 ± 0.004), AUC (0.994 ± 0.002) and specificity (0.989 ± 0.003). Like COVID-19 dataset, CNN-based model performed better, such as ResNet and DAX_Net. In Fig. 5 (c), we can intuitively observe the outstanding performance and robust stability of TetraFuse, while also noting its potential for generalization across diverse tasks.

**Table 6.** Model performance comparison on Kvasir dataset [68] with top-3 ranking markers: single underline (1st), double underline (2nd), <u>wavy underline</u> (3rd). All ranked values are bolded.

| Model Name | Acc | F1 | MCC | Kappa | AUC | Specificity |
| --- | --- | --- | --- | --- | --- | --- |
| beit_base [40] | 0.656 ± 0.020 | 0.644 ± 0.022 | 0.612 ± 0.023 | 0.608 ± 0.023 | 0.957 ± 0.006 | 0.951 ± 0.003 |
| gcvit_small [41] | 0.583 ± 0.057 | 0.564 ± 0.065 | 0.530 ± 0.064 | 0.524 ± 0.065 | 0.936 ± 0.015 | 0.940 ± 0.008 |
| convit_base [42] | 0.602 ± 0.007 | 0.563 ± 0.025 | 0.556 ± 0.009 | 0.545 ± 0.008 | 0.945 ± 0.004 | 0.943 ± 0.001 |
| efficientvit [43] | 0.734 ± 0.030 | 0.711 ± 0.069 | 0.701 ± 0.040 | 0.696 ± 0.049 | 0.962 ± 0.033 | 0.962 ± 0.019 |
| efficientformer [11] | 0.593 ± 0.066 | 0.572 ± 0.069 | 0.542 ± 0.074 | 0.534 ± 0.075 | 0.938 ± 0.020 | 0.942 ± 0.009 |
| davit_small [44] | 0.638 ± 0.004 | 0.627 ± 0.011 | 0.593 ± 0.005 | 0.587 ± 0.005 | 0.951 ± 0.001 | 0.948 ± 0.000 |
| swin_tiny [45] | 0.578 ± 0.060 | 0.533 ± 0.114 | 0.531 ± 0.099 | 0.517 ± 0.103 | 0.933 ± 0.038 | 0.940 ± 0.013 |
| swin_small [45] | 0.658 ± 0.028 | 0.641 ± 0.047 | 0.615 ± 0.029 | 0.609 ± 0.032 | 0.958 ± 0.003 | 0.951 ± 0.004 |
| swin_base [45] | 0.628 ± 0.046 | 0.594 ± 0.069 | 0.587 ± 0.050 | 0.575 ± 0.053 | 0.954 ± 0.011 | 0.947 ± 0.006 |
| visformer_small [46] | 0.605 ± 0.034 | 0.574 ± 0.054 | 0.558 ± 0.035 | 0.548 ± 0.039 | 0.917 ± 0.023 | 0.944 ± 0.005 |
| vit_small [25] | 0.613 ± 0.011 | 0.556 ± 0.032 | 0.573 ± 0.009 | 0.558 ± 0.012 | 0.953 ± 0.002 | 0.945 ± 0.002 |
| convformer_s36 [47] | 0.723 ± 0.089 | 0.706 ± 0.111 | 0.688 ± 0.097 | 0.683 ± 0.102 | 0.968 ± 0.015 | 0.960 ± 0.013 |
| mixer_b16_224 [48] | 0.763 ± 0.006 | 0.763 ± 0.006 | 0.730 ± 0.007 | 0.729 ± 0.016 | 0.974 ± 0.001 | 0.966 ± 0.001 |
| densenet161 [23] | 0.875 ± 0.019 | 0.876 ± 0.019 | 0.858 ± 0.022 | 0.858 ± 0.022 | 0.990 ± 0.002 | 0.982 ± 0.003 |
| resnet50 [49] | 0.870 ± 0.016 | 0.870 ± 0.016 | 0.852 ± 0.018 | 0.851 ± 0.019 | 0.990 ± 0.001 | 0.982 ± 0.002 |
| convnext_tiny [50] | 0.843 ± 0.006 | 0.843 ± 0.005 | 0.821 ± 0.006 | 0.821 ± 0.006 | 0.986 ± 0.001 | 0.977 ± 0.001 |
| convnext_small [50] | 0.835 ± 0.007 | 0.834 ± 0.007 | 0.811 ± 0.008 | 0.811 ± 0.008 | 0.986 ± 0.001 | 0.976 ± 0.001 |
| convnext_base [50] | 0.840 ± 0.002 | 0.840 ± 0.002 | 0.818 ± 0.003 | 0.817 ± 0.003 | 0.986 ± 0.001 | 0.977 ± 0.000 |
| inception_v4 [4] | 0.738 ± 0.020 | 0.732 ± 0.021 | 0.703 ± 0.023 | 0.701 ± 0.023 | 0.969 ± 0.004 | 0.962 ± 0.003 |
| HiFuse_Tiny [15] | 0.815 ± 0.046 | 0.813 ± 0.047 | 0.790 ± 0.051 | 0.789 ± 0.052 | 0.981 ± 0.006 | 0.974 ± 0.007 |
| HiFuse_Small [15] | 0.768 ± 0.069 | 0.765 ± 0.071 | 0.737 ± 0.076 | 0.735 ± 0.078 | 0.975 ± 0.011 | 0.967 ± 0.010 |
| HiFuse_Base [15] | 0.729 ± 0.054 | 0.721 ± 0.056 | 0.695 ± 0.060 | 0.690 ± 0.062 | 0.969 ± 0.009 | 0.961 ± 0.008 |
| DAX_Net [51] | 0.882 ± 0.028 | 0.881 ± 0.029 | 0.865 ± 0.033 | 0.865 ± 0.033 | 0.991 ± 0.003 | 0.983 ± 0.001 |
| Mamba_Tiny [21] | 0.708 ± 0.061 | 0.707 ± 0.062 | 0.668 ± 0.070 | 0.666 ± 0.070 | 0.963 ± 0.011 | 0.958 ± 0.009 |
| Mamba_Small [21] | 0.727 ± 0.066 | 0.722 ± 0.068 | 0.692 ± 0.073 | 0.689 ± 0.075 | 0.967 ± 0.012 | 0.961 ± 0.009 |
| Mamba_Base [21] | 0.729 ± 0.065 | 0.725 ± 0.068 | 0.693 ± 0.072 | 0.690 ± 0.074 | 0.967 ± 0.013 | 0.961 ± 0.009 |
| DAFNet_Tiny [30] | 0.884 ± 0.005 | 0.883 ± 0.005 | 0.868 ± 0.006 | 0.867 ± 0.006 | 0.991 ± 0.000 | 0.983 ± 0.001 |
| DAFNet_Small [30] | 0.883 ± 0.005 | 0.883 ± 0.005 | 0.866 ± 0.005 | 0.866 ± 0.005 | 0.991 ± 0.001 | 0.983 ± 0.001 |
| DAFNet_Base [30] | 0.888 ± 0.004 | 0.887 ± 0.005 | 0.873 ± 0.004 | 0.872 ± 0.005 | <u>0.992 ± 0.000</u> | 0.984 ± 0.001 |
| DAFNet_Large [30] | 0.881 ± 0.003 | 0.881 ± 0.004 | 0.865 ± 0.004 | 0.865 ± 0.004 | 0.990 ± 0.001 | 0.983 ± 0.001 |
| **TetraFuse_Tiny** | **<u>0.913 ± 0.003</u>** | **<u>0.913 ± 0.005</u>** | **<u>0.901 ± 0.004</u>** | **<u>0.901 ± 0.006</u>** | **<u>0.992 ± 0.003</u>** | **<u>0.987 ± 0.002</u>** |
| **TetraFuse_Small** | **<u>0.917 ± 0.004</u>** | **<u>0.916 ± 0.004</u>** | **<u>0.905 ± 0.005</u>** | **<u>0.905 ± 0.005</u>** | **<u>0.994 ± 0.001</u>** | **<u>0.988 ± 0.001</u>** |
| **TetraFuse_Base** | **<u>0.926 ± 0.006</u>** | **<u>0.925 ± 0.005</u>** | **<u>0.915 ± 0.004</u>** | **<u>0.915 ± 0.004</u>** | **<u>0.994 ± 0.002</u>** | **<u>0.989 ± 0.003</u>** |

#### 4.4.4. Comparative analysis of results

TetraFuse across three benchmarks—thoracic (COVID-19), dermatoscopic (ISIC 2018), and endoscopic—stems (Kvasir) from three architectural pillars:

- **Local-Global Synergy:** By combining CNN inductive bias with the CCDA module, TetraFuse captures both fine-grained pathological textures and global anatomical context.
- **Statistical Noise Filtering:** The LVGE enhancer uses second-order variance as a pathological filter to suppress early-stage artifacts (e.g., reflections or sensor noise), ensuring predictive stability.
- **Frequency-Aware Scaling:** Heteroscale kernels and the HFBI mechanism preserve sharp boundaries and multi-scale details that are typically lost during sequential downsampling.

In conclusion, by integrating spatial, channel, statistical, and frequency domains, TetraFuse achieves a state-of-the-art balance between computational efficiency and diagnostic precision.

#### 4.4.5. Comparison analysis of model efficiency

Table 7 compares the floating-point operations (FLOPs) and number of parameters across all the datasets. In terms of computational complexity and parameter efficiency, the TetraFuse models (ranging from 0.35 to 1.40 GFLOPs) demonstrate significant progress. Experimental results on model efficiency underscore the architectural superiority of TetraFuse. Compared to the classic ResNet50, Tetra-Fuse_Tiny achieves a transformative 91.53% reduction in FLOPs, proving that our omni-dimensional fusion strategy can distill essential pathological features with significantly fewer operations.

**Table 7.**
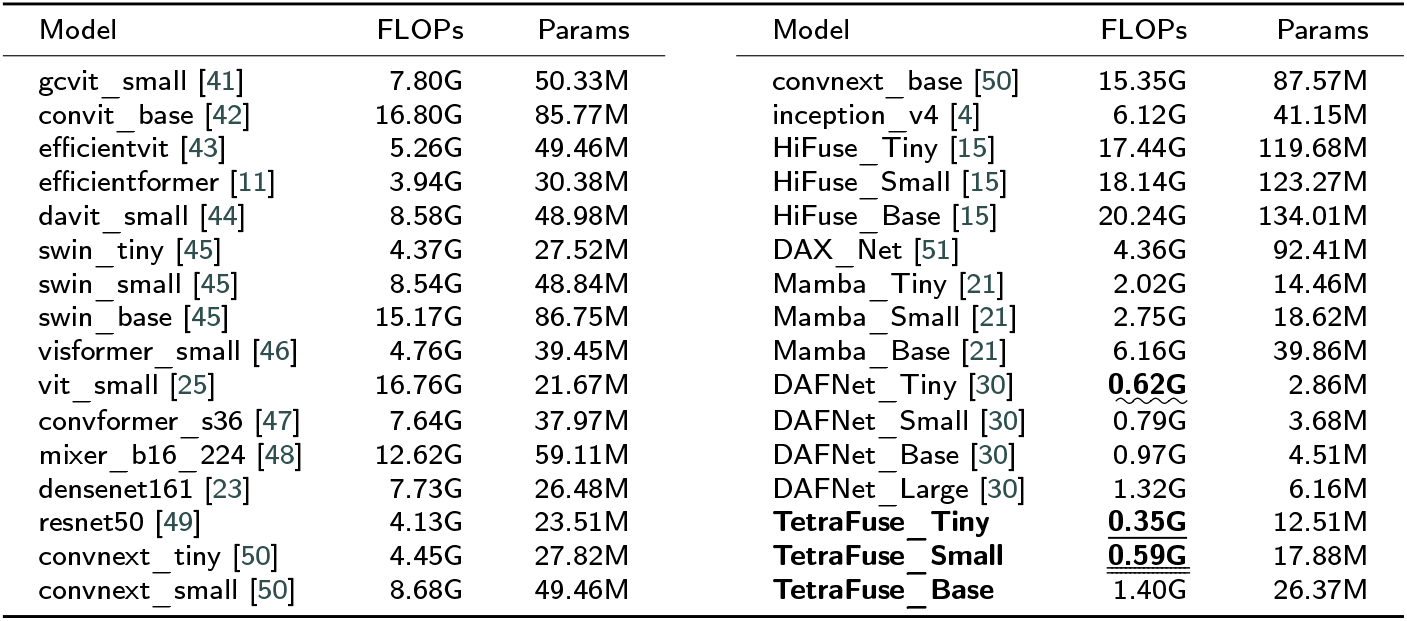
Model efficiency comparison: FLOPs (G) and Params (M). Our TetraFuse variants demonstrate superior efficiency compared to established vision backbones. Top-3 markers: single underline (1st), double underline (2nd), <u>wavy underline</u> (3rd). All ranked values are bolded.

Furthermore, even when pitted against the Mamba-based architectures (e.g., MedMamba), which are renowned for their linear complexity, TetraFuse_Tiny maintains a substantial lead with an 82.67% lower computational overhead. It is noteworthy that while TetraFuse_Tiny (12.51M) has a larger parameter footprint compared to ultra-compact models such as DAFNet_Tiny (2.86M), this is a strategic design trade-off. The additional parameters are primarily attributed to the weight-generation sub-networks within the CCDA and LFA modules, which are essential for generating dynamic kernels that adapt to complex pathological textures.

From a deployment perspective, although parameters dictate the memory storage requirement, FLOPs are typically the primary bottleneck for efficient inference latency on GPU servers and modern AI-accelerated edge devices. Given that 12.51M parameters only occupy approximately 50MB of memory (in FP16 precision), which is well within the capacity of most mobile medical hardware, TetraFuse prioritizes minimizing FLOPs and maximizing diagnostic accuracy over extreme parameter compression. This suggests that by synergizing spatial, channel, and frequency-domain information, TetraFuse avoids the heavy sequential scanning costs inherent in Mamba-like models while delivering superior diagnostic precision. Such efficiency makes TetraFuse an ideal candidate for efficient clinical deployment on resource-constrained mobile medical devices. Fig 6 provides a clearer visual comparison of the results.

**Figure 6.**
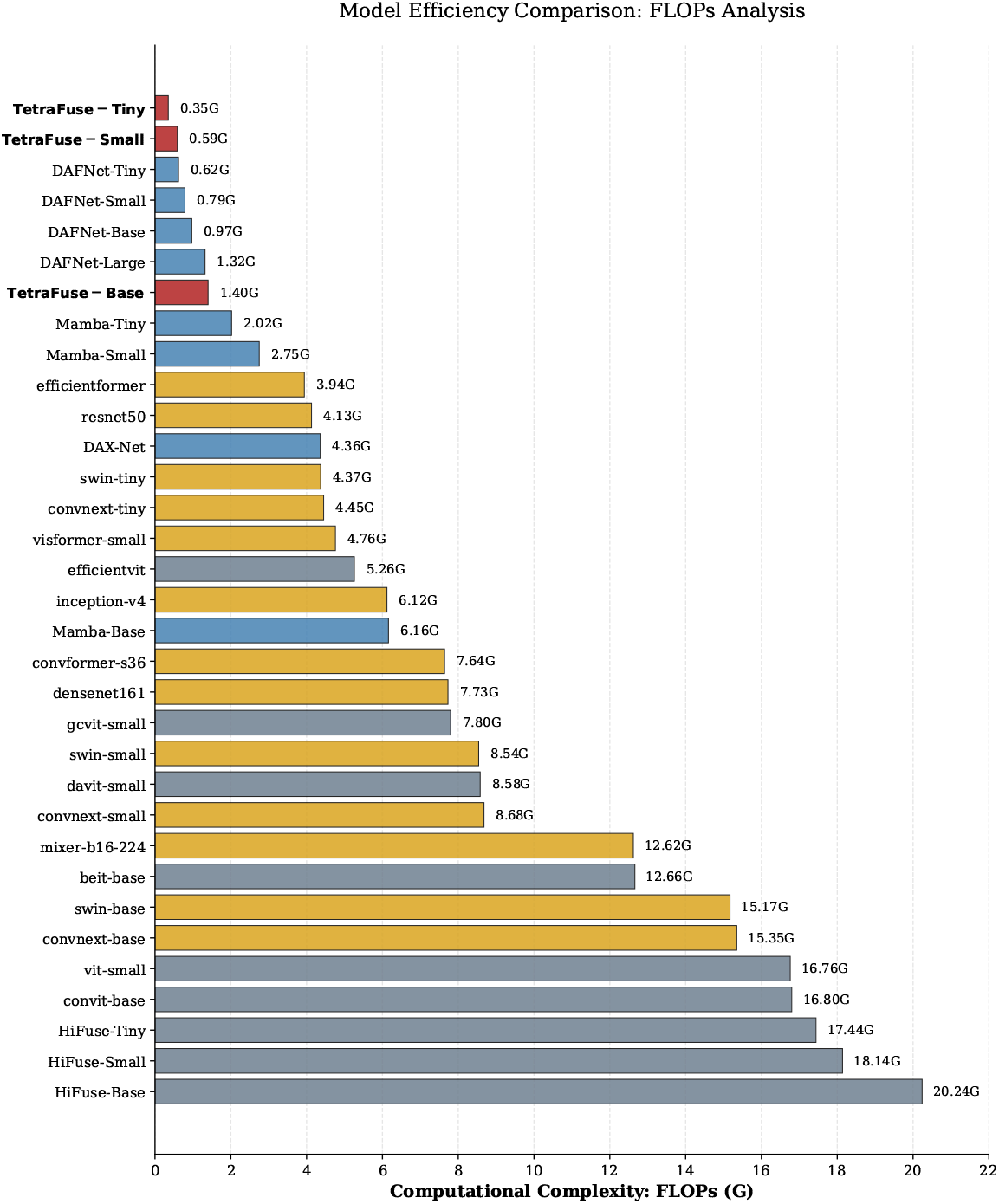
FLOPs results on different models.

#### 4.4.6. Confusion matrix and ROC curve

Fig 7 presents the corresponding matrices and ROC curves of different models. From the confusion matrix analysis, all models performed better on COVID-19 than other datasets. Among them, TetraFuse_Base achieved the most balanced and best classification performance overall. On the ISIC 2018 dataset, other models struggled with the intraepithelial carcinoma, melanoma and actinic keratoses categories; however, our TetraFuse attained higher accuracy in these classes, and get best performance on other categories at the same time. For the Kvasir dataset, the models commonly classified the normal z-line category as esophagitis by mistake. Nonetheless, the TetraFuse attained the most consistent and superior results.

**Figure 7.**
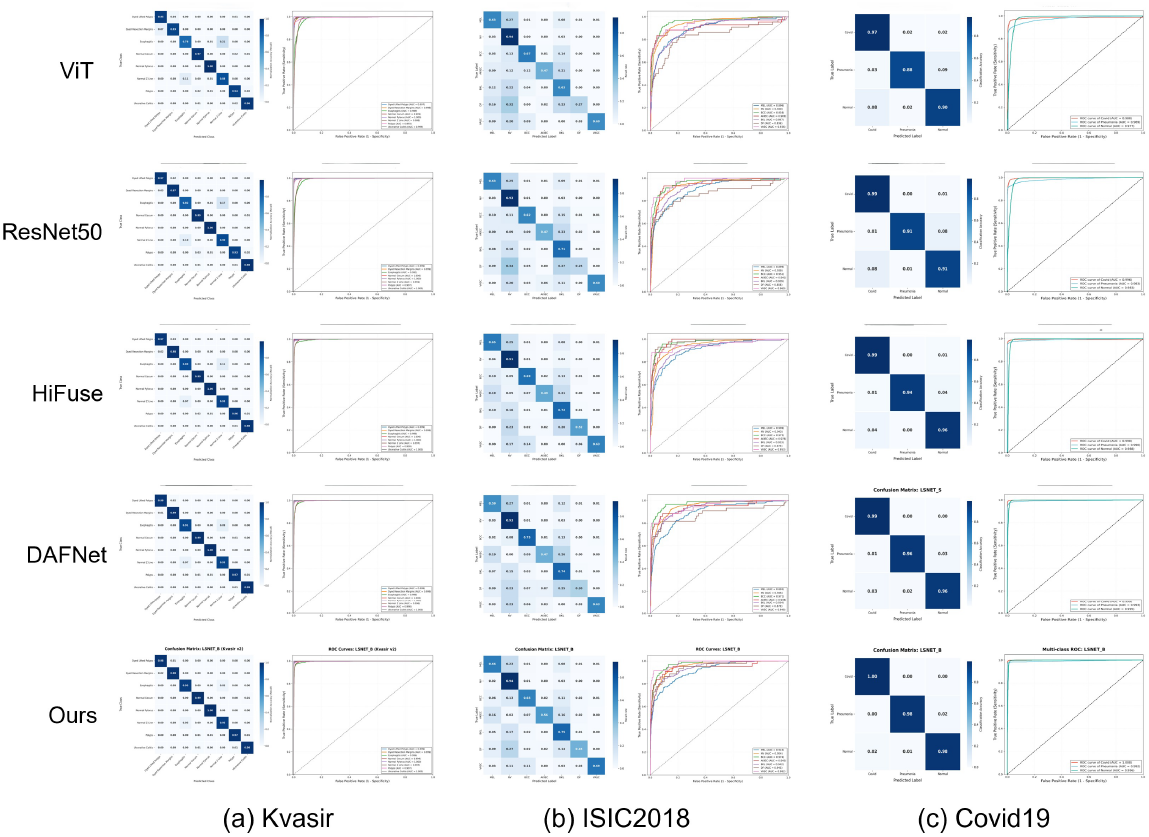
Confusion matrix and ROC curve of different models on three datasets.

### 4.5. Ablation Study

Totally we can find the clear contrast in Fig 8. To systematically evaluate the individual contribution and the synergistic effect of the proposed modules within the **TetraFuse** framework, we conduct incremental ablation experiments on three heterogeneous medical datasets. As summarized in Table 8, the original baseline refers to a simplified backbone architecture derived from TetraFuse by removing all proposed fusion modules, including CCDA, LVGE, and HFBI. Specifically, all CCDA blocks are replaced with standard depthwise separable convolutions, and no statistical or frequency-domain enhancement is applied. This ensures that the overall macro-architecture (MSC-Stem, stage hierarchy, and classifier) remains unchanged, enabling fair attribution of performance gains. Each component—namely *Heteroscale Selective Perception* (HSP), *Local Feature Aggregation* (LFA), *High-Frequency Boundary Injector* (HFBI), and *Local Variance Guided Enhancer* (LVGE)—is progressively integrated into the network.

**Table 8.** Component ablation experiment results across datasets. HSP means heteroscale selective perception, LFA means local feature aggregation, LVGE means local variance guided enhancer, and HFBI means high-frequency boundary injector.

| COVID-19 dataset |  |  |  |  |  |  |
| --- | --- | --- | --- | --- | --- | --- |
| Component | Acc | F1 | MCC | Kappa | AUC | Specificity |
| original | 0.913 ± 0.018 | 0.921 ± 0.018 | 0.891 ± 0.017 | 0.888 ± 0.017 | 0.960 ± 0.005 | 0.946 ± 0.009 |
| + HSP | 0.924 ± 0.015 | 0.931 ± 0.015 | 0.903 ± 0.021 | 0.900 ± 0.022 | 0.969 ± 0.007 | 0.950 ± 0.007 |
| + LFA | 0.935 ± 0.018 | 0.940 ± 0.015 | 0.913 ± 0.015 | 0.912 ± 0.012 | 0.975 ± 0.010 | 0.959 ± 0.006 |
| + HFBI | 0.950 ± 0.012 | 0.949 ± 0.010 | 0.926 ± 0.015 | 0.925 ± 0.014 | 0.984 ± 0.006 | 0.971 ± 0.007 |
| + LVGE | **0.956 ± 0.003** | **0.956 ± 0.005** | **0.935 ± 0.008** | **0.934 ± 0.008** | **0.990 ± 0.005** | **0.979 ± 0.004** |
| ISIC 2018 dataset |  |  |  |  |  |  |
| Component | Acc | F1 | MCC | Kappa | AUC | Specificity |
| original | 0.801 ± 0.006 | 0.676 ± 0.016 | 0.671 ± 0.011 | 0.645 ± 0.011 | 0.947 ± 0.018 | 0.949 ± 0.003 |
| + HSP | 0.815 ± 0.014 | 0.688 ± 0.019 | 0.685 ± 0.011 | 0.649 ± 0.010 | 0.955 ± 0.015 | 0.951 ± 0.003 |
| + LFA | 0.824 ± 0.008 | 0.699 ± 0.010 | 0.694 ± 0.010 | 0.658 ± 0.010 | 0.960 ± 0.014 | 0.954 ± 0.001 |
| + HFBI | 0.830 ± 0.010 | 0.710 ± 0.013 | 0.703 ± 0.021 | 0.668 ± 0.022 | 0.962 ± 0.002 | 0.959 ± 0.005 |
| + LVGE | **0.835 ± 0.009** | **0.722 ± 0.013** | **0.710 ± 0.011** | **0.710 ± 0.012** | **0.966 ± 0.011** | **0.961 ± 0.002** |
| Kvasir dataset |  |  |  |  |  |  |
| Component | Acc | F1 | MCC | Kappa | AUC | Specificity |
| original | 0.899 ± 0.009 | 0.897 ± 0.010 | 0.882 ± 0.010 | 0.885 ± 0.011 | 0.980 ± 0.001 | 0.979 ± 0.001 |
| + HSP | 0.905 ± 0.007 | 0.904 ± 0.006 | 0.891 ± 0.007 | 0.892 ± 0.007 | 0.986 ± 0.000 | 0.984 ± 0.001 |
| + LFA | 0.913 ± 0.009 | 0.912 ± 0.009 | 0.900 ± 0.011 | 0.900 ± 0.009 | 0.990 ± 0.001 | 0.987 ± 0.001 |
| + HFBI | 0.920 ± 0.006 | 0.920 ± 0.007 | 0.910 ± 0.007 | 0.910 ± 0.007 | 0.993 ± 0.001 | 0.988 ± 0.001 |
| + LVGE | **0.926 ± 0.006** | **0.925 ± 0.005** | **0.915 ± 0.004** | **0.915 ± 0.004** | **0.994 ± 0.002** | **0.989 ± 0.003** |

**Figure 8.**
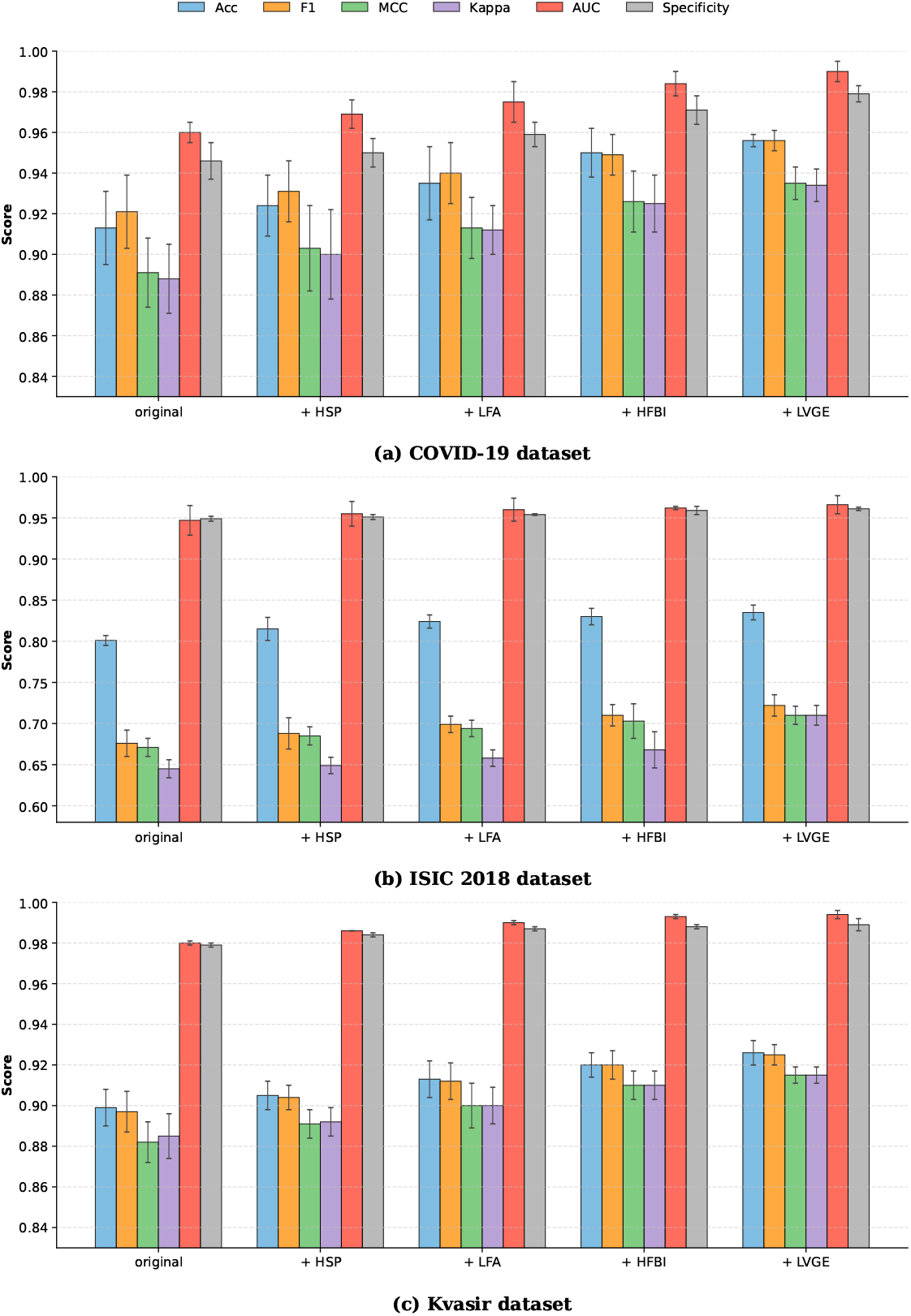
Component-wise Ablation Study of TetraFuse: Performance Comparison Shown by Histogram.

#### 4.5.1. Impact of Spatial Domain Optimization (HSP & LFA)

The integration of HSP yields a consistent performance gain; conversely, removing HSP from the full TetraFuse configuration leads to a significant drop in Accuracy (Acc) on the ISIC 2018 dataset from 0.835 back towards 0.815, demonstrating that the multi-scale perception mechanism effectively handles the significant variability in lesion sizes. Furthermore, the LFA module strengthens the localized inductive bias; as evidenced in the COVID-19 dataset, its addition boosts Acc to 0.935, confirming that aggregating localized spatial features is vital for detecting subtle pathological textures.

#### 4.5.2. Significance of Frequency Domain Injection (HFBI)

Introducing HFBI allows the network to re-acquire high-frequency boundary cues that are often attenuated during sequential downsampling. This is clearly reflected in the Kvasir dataset, where the AUC score improves to 0.993. By re-injecting sharp frequency components, TetraFuse maintains structural integrity and boundary definition, which is critical for distinguishing polyps from surrounding healthy tissue in endoscopic imagery.

#### 4.5.3. Statistical Refinement and Artifact Suppression (LVGE)

The full TetraFuse model, attained with the inclusion of the LVGE module, achieves the most superior performance. On the COVID-19 benchmark, the Specificity reaches 0.979, while the F1-score on ISIC 2018 reaches 0.722 (a 6.8% relative improvement over the Original baseline). LVGE’s utilization of second-order statistical variance acts as a pathological filter, effectively suppressing non-informative artifacts such as specular reflections or sensor noise, thereby ensuring the stability and reliability of the diagnostic predictions.

#### 4.5.4. Overall Synergy

The incremental performance trajectory shown in Table 8 underscores the synergistic nature of the Tetra-domain strategy. Each module addresses a specific limitation spatial scale, locality, frequency definition, or statistical noise enabling TetraFuse to construct a robust and comprehensive feature representation for complex clinical tasks. By systematically removing one module from the full **TetraFuse** framework, we observe a consistent performance degradation (Δ) across all benchmarks. Specifically, the exclusion of **HFBI** leads to the most substantial drop in Accuracy on the COVID-19 dataset (−1.5%), reinforcing its role in recovering critical boundary information. Similarly, the removal of **HSP** on the ISIC 2018 dataset results in a 1.4% decrease, confirming its necessity for multi-scale lesion perception. Our sensitivity analysis also indicates that while the integration sequence can vary, the spatial-to-frequency pipeline (HSP → LFA → HFBI → LVGE) provides the most robust convergence. This Leave-one-out evidence confirms that no module is redundant; instead, they form a functionally complementary architecture for complex medical diagnostic tasks. The details are shown in Table 9. In addition, the improvement in classification ability of the architecture for three datasets is clearly shown in the following Fig 9 as a t-SNE plot.

**Table 9.** Leave-one-out necessity analysis of TetraFuse across three datasets. “Δ” denotes the performance drop in Accuracy (Acc) when a specific module is removed from the full framework.

| Configuration | COVID-19 |  | ISIC 2018 |  | Kvasir |  |
| --- | --- | --- | --- | --- | --- | --- |
| | Acc | $\Delta$ | Acc | $\Delta$ | Acc | $\Delta$ |
| **Full TetraFuse** | **0.956** | - | **0.835** | - | **0.926** | - |
| w/o LVGE | 0.950 | -0.6% | 0.830 | -0.5% | 0.920 | -0.6% |
| w/o HFBI | 0.941 | -1.5% | 0.829 | -0.6% | 0.919 | -0.7% |
| w/o LFA | 0.945 | -1.1% | 0.826 | -0.9% | 0.918 | -0.8% |
| w/o HSP | 0.945 | -1.1% | 0.821 | -1.4% | 0.920 | -0.6% |
| Baseline | 0.913 | -4.3% | 0.801 | -3.4% | 0.899 | -2.7% |

**Figure 9.**
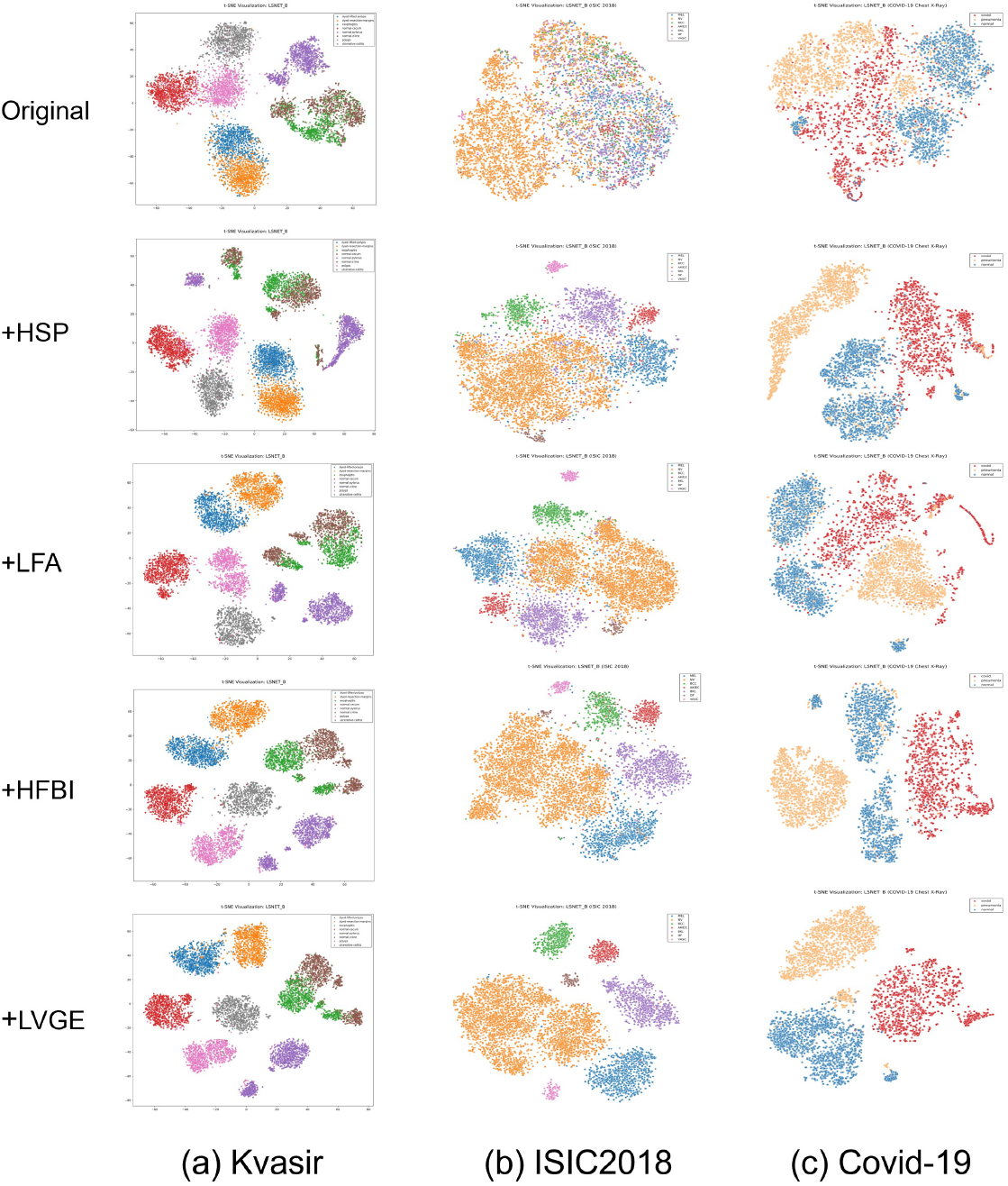
t-SNE visualizations on three datasets.

### 4.6. Visual Inspection

To validate the clinical interpretability of TetraFuse, we visualize the attention maps using Grad-CAM across three heterogeneous datasets in Fig 10.

**Figure 10.**
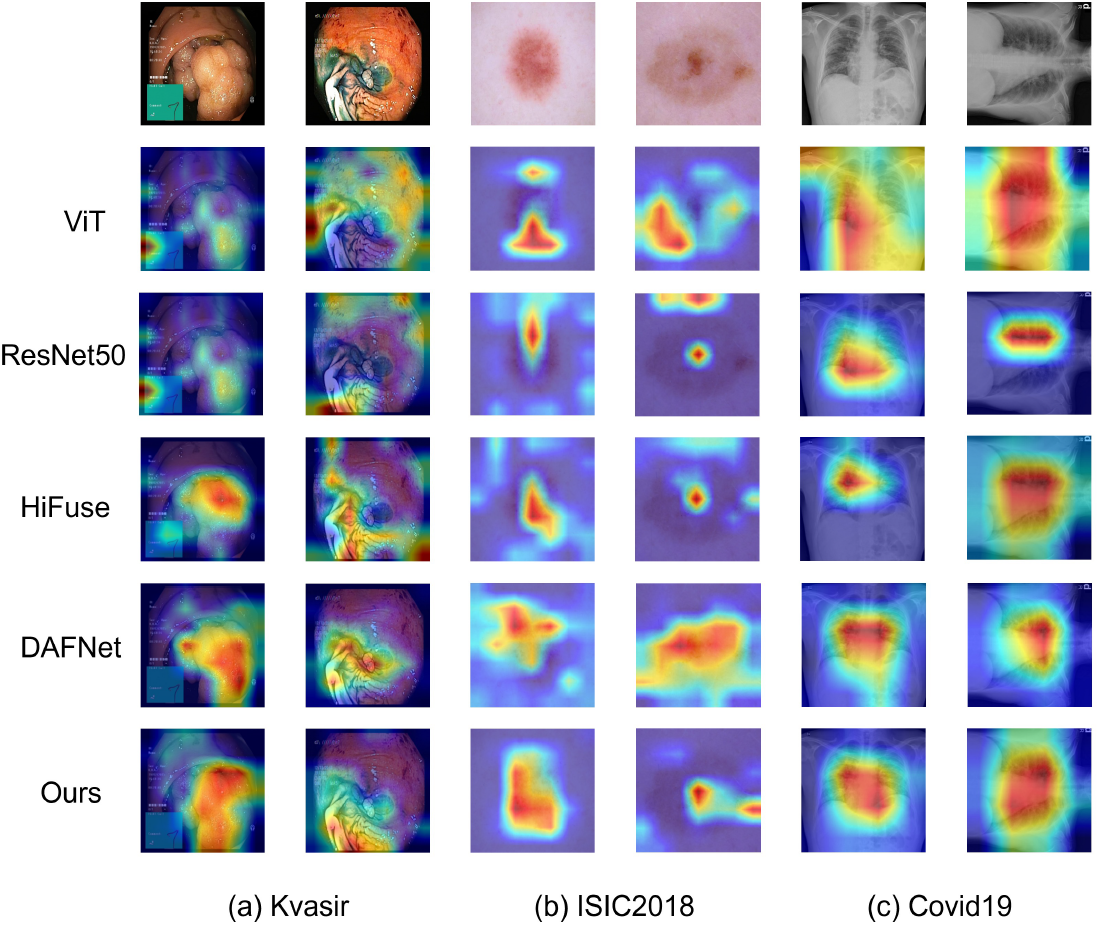
Grad-CAM visualizations on three datasets.

Artifact Suppression (Kvasir): While baselines are often distracted by specular reflections and mucosal noise, TetraFuse maintains a concentrated focus on polyp regions. This is enabled by the LVGE module, which utilizes secondorder statistics to filter out non-pathological optical artifacts. Boundary Delineation (ISIC 2018): For skin lesions with blurred edges, TetraFuse provides significantly sharper heatmaps than DAFNet. The HFBI module re-injects highfrequency boundary cues, ensuring the model precisely aligns with the pathological contours. Scale Adaptation (COVID-19): In thoracic X-rays, TetraFuse accurately localizes subtle pneumonia patches of varying sizes without attention drift to skeletal structures. This stems from the ability of the HSP module to adaptively adjust the receptive field to match localized textural heterogeneities.

In general, the high spatial alignment between TetraFuse’s activations and ground-truth lesions confirms its diagnostic reliability and consistency with clinical prior knowledge.

## 5. Discussion and Limitations

### 5.1. Mechanisms of Omni-Dimensional Fusion

The robust performance of TetraFuse across three heterogeneous medical datasets—thoracic radiography, dermatoscopy, and endoscopy—demonstrates a state-of-the-art balance between diagnostic precision and computational economy. We attribute this success to the synergistic integration of inductive bias and global context. While Transformer-based architectures often require massive pretraining to stabilize macroscopic anatomical context, the convolutional-heavy backbone of TetraFuse naturally captures localized textural heterogeneities, such as pneumonia patches in COVID-19 radiographs.

Furthermore, TetraFuse transcends the limitations of existing attention paradigms. While mainstream methods like Coordinate Attention [59] or Triplet Attention [60] are primarily confined to 2D spatial-channel interactions, our framework introduces a 4D synergy. By embedding statistical noise-filtering (LVGE) and high-frequency boundary injection (HFBI) directly into the feature hierarchy, TetraFuse ensures that deep semantic features remain both clean and sharp, which is reflected in the significantly improved AUC and Kappa scores observed in the Kvasir and ISIC experiments.

### 5.2. Architectural Evolution and Efficiency

A critical comparison with its predecessor, DAFNet [30], reveals a significant evolution in computational scalability. DAFNet relies on heavy attention maps that scale poorly with image resolution. In contrast, TetraFuse utilizes the CCDA module, which employs a parameter-free shuffle mechanism to facilitate cross-group synergy. By reconstructing global topological dependencies without redundant parameters, TetraFuse successfully eliminates information isolation.

The experimental results highlight this extreme efficiency: compared to the classic ResNet50 and the state-of-the-art Mamba-Tiny, TetraFuse-Tiny reduces FLOPs by 91.53% and 82.67%, respectively, without sacrificing accuracy. This favorable performance-to-parameter ratio positions TetraFuse as a deployment-ready paradigm for resource-constrained mobile medical devices.

### 5.3. Operational Boundaries and Error Analysis

Deep analysis of the operational boundaries of TetraFuse yields critical insights into its design choices. The reliance on second-order statistics within the LVGE module is highly effective when the signal-to-noise ratio (SNR) is relatively high and noise follows a near-Gaussian distribution. However, its efficacy may diminish in scenarios characterized by extremely low-contrast textures where pathological signals are indistinguishable from sensor noise. Consequently, we restrict LVGE to the initial stages, transitioning to frequency-domain injection (HFBI) in deeper layers to capture abstract semantic boundaries.

This mechanistic boundary is further reflected in our visual inspection of misclassified samples (Fig. 11). In the ISIC 2018 dataset, errors primarily occurred between Melanocytic Nevus and Seborrheic Keratoses, typically involving: (1) *Extremely fuzzy boundaries* where the lesion texture seamlessly blends into healthy skin, and (2) *Physical occlusions* such as dense hair or oil artifacts. While the LVGE module filters standard statistical noise, it remains challenged by dense occlusions that physically block pathological signals. Additionally, the slight performance gap of TetraFuse-Tiny compared to DAFNet on ISIC 2018 suggests that in tasks with massive intra-class variance, a certain level of structural redundancy (as seen in DAFNet) might be beneficial for capturing outlier features.

**Figure 11.**
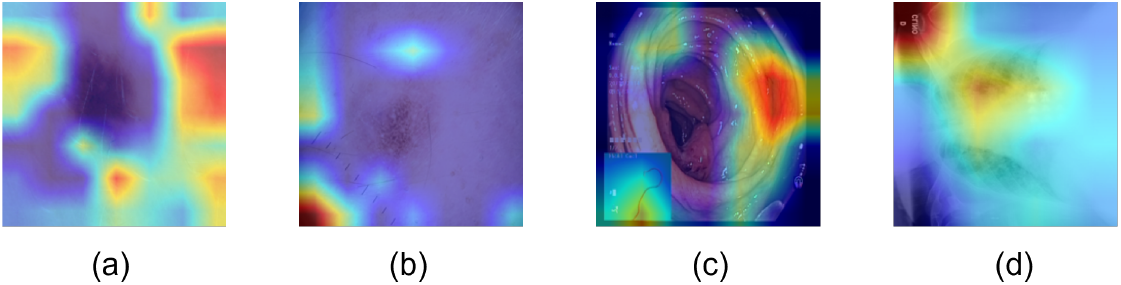
four selected misclassified samples.

### 5.4. Limitations and Future Work

Despite its promising results, several limitations of TetraFuse warrant further investigation:

- **Volumetric Modality Extension:** Currently, the omni-dimensional fusion logic is optimized for 2D imagery. Future work will focus on extending these kernels into 3D spaces to capture spatio-temporal dependencies in volumetric data such as CT and MRI stacks.
- **Long-tail Distribution:** Consistent with most deep learning models, TetraFuse’s sensitivity dips when dealing with extremely rare pathological classes. Incorporating Knowledge Graph Filtering or clinical prior biases into the fusion block could potentially enhance representation for long-tail samples.
- **Clinical Explainability:** While Grad-CAM visualizations demonstrate that TetraFuse localizes correct lesion areas, a more rigorous interpretability framework is required. We aim to integrate uncertainty estimation or fuzzy logic into the fusion process to provide clinicians with robust confidence intervals for diagnostic trust.
- **Trade-off Between Parameters and Dynamic Expression:** Although TetraFuse_Tiny achieves superior FLOPs efficiency, its parameter footprint (12.51M) is larger than some ultra-compact static models like DAFNet_Tiny. This overhead is primarily localized in the weight-generation sub-networks required for dynamic kernel synthesis. Future research will explore model compression techniques, such as parameter pruning or knowledge distillation, to further condense these auxiliary networks without sacrificing the adaptive representational power of the omni-dimensional fusion mechanism.

## 6. Conclusion

In this paper, we presented TetraFuse, a novel and highly efficient framework tailored for multi-modal medical image classification. To address the inherent challenges of information isolation, pathological noise, and boundary degradation in clinical imagery, we proposed a synergistic fusion strategy across four critical dimensions: *Space, Channel, Statistics, and Frequency*.

Specifically, the CCDA module reconstructs global topological dependencies to eliminate channel redundancy, while the LVGE enhancer leverages second-order statistical variance to suppress early-stage artifacts. Furthermore, the integration of HSP and HFBI ensures multi-scale adaptability and sharp boundary retention, effectively resolving the over-smoothing issue in deep semantic features.

Extensive experiments on three diverse medical datasets (COVID-19, ISIC 2018, and Kvasir) demonstrate that TetraFuse significantly outperforms state-of-the-art CNN and Transformer-based models. By delivering a state-of-the-art balance between computational economy and predictive stability, TetraFuse establishes a highly dependable paradigm for efficient clinical diagnostics on resource-constrained devices.

## CRediT authorship contribution statement

**Yufei Gao:** Conceptualization, Methodology, Software, Formal analysis. **Jiaqi Li:** Methodology, Software, Validation, Writing - original draft, Visualization **Jing Xu:** Software, Data curation, Visualization. **Qing Li:** Validation, Investigation, Visualization. **Ziyu Li:** Investigation, Formal analysis. **Guohua Zhao:** Resources, Validation. **Yucheng Shi:** Resources, Data curation. **Xia Wu:** Conceptualization, Funding acquisition, Project administration, Supervision, Writing - review & editing. **Yameng Zhang:** Conceptualization, Resources, Supervision, Writing - review & editing.

## Acknowledgments

This work was supported in part by the National Science Fund for Distinguished Young Scholars of China (62325601), the National Natural Science Foundation of China (92470125, 62506037, 82402395), Natural Science Foundation of Henan Province (262300421812), Fundamental Research Funds for the Central Universities (XSQD-6120240006), and Henan Province Medical Science and Technology Research Co-Development Program (LHGJ20250037).

